# NPTX2-Centered Cognitive Resilience Mechanisms in the Context of AD Pathology

**DOI:** 10.1101/2025.10.17.683150

**Authors:** Yuelin Lao, Mei-Fang Xiao, Shiyu Ji, Ignazio S. Piras, Kyungdo Kim, Anna Bonfitto, Serena Song, Arailym Aldabergenova, Chan-Hyun Na, Jennifer Sloan, Angela Trejo, Changiz Geula, Emily J. Rogalski, Claudia H. Kawas, Maria M. Corrada, Geidy E. Serrano, Thomas G. Beach, Juan C. Troncoso, Matthew J. Huentelman, Carol A. Barnes, Paul F. Worley, Carlo Colantuoni

**Author notes:** Lead contact and correspondence (P.F.W.) & (C.C.).

## Abstract

**Background:** Cognitive resilience to Alzheimer’s disease (AD) pathology is associated with preserved expression of NPTX2, an activity-regulated synaptic protein involved in circuit plasticity, excitation–inhibition balance, and complement-linked synapse regulation. However, the broader molecular programs coordinated with NPTX2 in resilient individuals remain unclear.

**Methods:** We analyzed postmortem middle temporal gyrus tissue using targeted PRM-MS proteomics in 135 individuals and bulk RNA-seq in an expanded 575-sample cohort. NPTX2-associated molecular coordination was assessed within cognitively normal low-pathology controls (CN-Lo), cognitively normal high-pathology controls (CN-Hi), mild cognitive impairment (MCI), and AD. Correlation-based approaches were applied using NPTX2 protein and NPTX2 mRNA expression as anchors to define resilience mechanisms in CN-Hi subjects.

**Results:** NPTX2 protein abundance was preserved across all controls regardless of age and pathology but reduced in MCI and AD. NPTX2 mRNA expression was also invariant across pathology within controls and reduced in MCI and AD but decreased markedly with age. Targeted proteomics identified NPTX2 relationships with synaptic and inhibitory-circuit proteins that were preserved across control groups, alongside CN-Hi-specific recruitment of trafficking, lysosomal, metabolic, and proteostasis-associated proteins. Transcriptome-wide correlations with NPTX2 revealed differences in gene co-expression between groups, identifying a prominent activity-dependent program including BDNF, VGF, SCG2, SST, SERTM1, DUSP4, and EGR4, that was preserved in both CN-Lo and CN-Hi subjects, while genes recruited to the NPTX2 network specifically in CN-Hi implicated immune, neuroprotective, translation, and proteostasis-related pathways. Coupling differential gene expression analysis with co-expression, we further identified five candidate resilience genes whose expression and NPTX correlation was preserved across controls, but lost in MCI and AD: SST, MAL2, TAC1, SERTM1, and RFK. Expression of genes in distinct NPTX2 co-expression classes can be freely explored in our bulk RNA-seq data and other public AD transcriptomic datasets at [NeMO Analytics](https://nemoanalytics.org/landing/neurodegeneration/).

**Conclusion:** Findings suggest that cognitive resilience in the context of AD neuropathology engages a coordinated molecular state distinct from both persevered cognition without pathology and MCI/AD, which is organized around preserved and selectively remodeled NPTX2-associations. Rather than reflecting broad transcript abundance changes, resilience was characterized by maintained synaptic and inhibitory programs, and adaptive proteostasis and trafficking pathways that distinguish resilient high-pathology individuals from low-pathology controls or symptomatic AD.

## Introduction

NPTX2 is an immediate early gene (IEG) and synaptic protein that plays a key role in memory consolidation by regulating the balance of excitation and inhibition within neuronal ensembles [1, 2]. In Alzheimer’s disease (AD), NPTX2 mRNA and protein levels are reduced in the postmortem neocortex compared to cognitively normal age-matched controls [3]. Notably, this NPTX2 reduction is not observed in certain individuals who remained cognitively intact until death, but who exhibited AD pathology upon postmortem examination [3]. This clinical-pathological condition is termed “asymptomatic AD” [4] or “high-pathology control” [5, 6].

Preservation of NPTX2 expression in these individuals first suggested its role as a resilience factor for preserved cognition. This hypothesis was further supported by a screen of postmortem brain tissue from aged individuals for glycoproteins linked to preserved cognition that identified multiple glycosylation states of NPTX2 as top candidates [7]. In cerebrospinal fluid (CSF), NPTX2 measures are reduced in individuals with mild cognitive impairment (MCI) and AD, and improve the diagnostic performance of established AD biomarkers including Aβ42/40 and tau [8]. Longitudinal studies further show that NPTX2 levels, together with tau measures, predict time to onset of cognitive symptoms [9]. Moreover, in a large unbiased CSF biomarker screen, the ratio of NPTX2/YWHAZ emerged as a highly effective diagnostic indicator [10].

Further evidence supporting a role for NPTX2 in cognitive resilience comes from rescue experiments in animal models where NPTX2 transgene expression restored synaptic and cognitive function in a rat model of age-related cognitive decline [11] and restored neuronal activity measures and visual processing in a mouse model of Aβ amyloidosis [12].

The “life cycle” of NPTX2 involves multiple cell-biological processes that can influence its expression. Its activity-dependent transcription and translation follow canonical IEG mechanisms [13], while its function as a glycosylated presynaptic protein requires proper Golgi processing, assembly into disulfide-linked complexes with NPTX1 and NPTXR, and fast axonal transport to presynaptic terminals [14]. NPTX2 additionally undergoes activity-dependent release at presynaptic terminals and enzymatic shedding during sleep-related circuit refinement [15].

Among the NPTX family members, NPTX2 is uniquely IEG-regulated and is essential for targeting the NPTX complex to excitatory synapses on parvalbumin (PV) interneurons [1]. At these sites, the transmembrane domain of NPTXR retains the NPTX1/2/R complex at the presynaptic membrane where it binds and recruits postsynaptic AMPA receptors, particularly GluA4 [2, 16]. Together with the activity dependent regulation of NPTX2, this synaptogenesis action enhances circuit-mediated inhibition and contributes to maintenance of excitation–inhibition balance within sparse, behaviorally engaged neuronal assemblies.

Another important known function of the NPTX1/2/R complex is inhibition of complement factor C1q [17]. C1q prominently associates with synapses and activates the classical complement cascade to promote microglial engulfment and synapse removal [17]. During sleep, C1q is normally released from NPTX1/2/R inhibition when the NPTXR transmembrane tether is cleaved by extracellular proteases, shedding the NPTX complex into the CSF [15]. This NPTX–C1q–microglial mechanism is relevant to physiological circuit refinement and potentially to pathological synapse loss in AD [17].

In the present study, we use NPTX2 as a point of reference for multiomic discovery of molecular correlates of resilience or transition to cognitive decline. Here, we examine NPTX2 mRNA and protein co-expression with 19 targeted proteins assayed by PRM-MS and the full transcriptome using bulk RNA-seq. We assessed NPTX2 co-expression within four clinical-pathological groups: low-pathology cognitively normal controls (CN-Lo), cognitively normal individuals with high AD pathology (CN-Hi), MCI, and AD. This identified transcripts with preserved, recruited, reversed, or suppressed, NPTX2 co-expression in CN-Hi compared with CN-Lo subjects, as well as relationships with NPTX2 that were pathology-disrupted (present in CN-Lo but absent in both CN-Hi and AD/MCI). Using differential expression analysis, we also identified candidate cognitive resilience genes whose expression is high in CN-Hi cases versus both CN-Lo and MCI/AD, indicating that some resilience mechanisms may be specifically engaged in the context of AD pathology (not simply preserved in both CN-Lo and CN-Hi).

## Methods

### Sample Acquisition

Fresh-frozen cortical punches from the middle temporal gyrus were obtained from four brain banks: Banner Sun Health Research Institute, University of California, Irvine, Johns Hopkins University, and the SuperAging Research Initiative.

### Brain Bank Contributors

#### Banner Sun Health Brain and Body Donation Program (BBDP)

The Brain and Body Donation Program at Banner Sun Health Research Institute (BSHRI) is a prospective clinicopathologic cohort initiated in 1987 that recruits mainly from retirement communities in metropolitan Phoenix, Arizona [18, 19]. Enrollees undergo standardized medical, neurologic, and neuropsychological assessments during life, with rapid autopsy at death (median post-mortem interval ∼3.0 hours) yielding high-quality frozen tissues, including a median brain RNA Integrity Number of 8.9 [18–21]. Whole-body donation has been available since 2005, and neuropathologic diagnoses are rendered by licensed pathologists using contemporary consensus criteria [18].

### 90+ Study

The 90+ Study is a population-based longitudinal cohort of adults aged 90 years and older, drawn primarily from surviving members of the Leisure World Cohort in Laguna Woods, Irvine, California [22]. Participants are evaluated approximately every 6 months with a neurologic examination, a standardized neuropsychological battery, and informant questionnaires; remote phone or mail assessments are used when in-person visits are not feasible [22, 23]. Dementia is diagnosed by study clinicians, and a brain-donation program provides neuropathologic confirmation using CERAD and NIA-Reagan criteria [22]. Additional cohort descriptions and epidemiologic findings on dementia prevalence and incidence in this population have been published [24, 25].

### SuperAging Research Initiative

This prospective, longitudinal cohort enrolls community-dwelling adults aged 80 years or older. SuperAgers are defined by performance at or above the mean for cognitively healthy 50–60-year-olds on Rey Auditory Verbal Learning Test delayed recall and at least average-for-age performance on non-memory measures; cognitively average older-adult controls perform within expected normative ranges for their demographic group on delayed recall and at least within the average range for age on non-memory measure [26].

Recruitment occurs through community-engaged research and collaborative outreach. Participants complete standardized longitudinal neuropsychological evaluations, with neuroimaging, biomarker studies, and optional brain donation for neuropathologic characterization. The cohort was expanded in 2021 as a multisite study with enrollment across five sites in the United States and Canada [27].

### Johns Hopkins Division of Neuropathology Brain Bank

This autopsy cohort draws brain donations through the Maryland Office of the Chief Medical Examiner and affiliated programs, explicitly including young adults; standardized dissections and region-specific sampling are performed at the Johns Hopkins Division of Neuropathology [28]. Tissues consisted of fresh-frozen punches of the left middle temporal gyrus. For each case, detailed medical and toxicology histories are abstracted, and tissue is processed for uniform histology, with genotyping and molecular assays performed when applicable [28]. The cohort has been used to characterize preclinical Alzheimer-type pathology in individuals aged 40–50, informing age-stratified analyses of early disease biology [28].

#### RNA-seq Workflow

FASTQ files were analyzed using the Nextflow RNA-seq pipeline [29], including trimming with Trim Galore, alignment with STAR [30], and quantification of counts at the gene level with SALMON [31]. Quality control was conducted using MultiQC [32] and Qualimap [33]. Samples were included if they had all covariates, at least 20 million sequencing reads, and if 80% of those reads were uniquely mapped to the human transcriptome. Raw counts were imported from DESeq2 [34] and, for quality control purposes, transformed using the vst method. We removed genes located on the sex chromosomes and conducted Principal Component Analysis (PCA) to identify outlier samples, defined as those that were above ± 3 standard deviations from the average of at least one of the two top PCs.

After quality control, a total of 575 samples were included in the analysis. The control group comprised 180 cognitively normal individuals aged over 50 years. The mild cognitive impairment (MCI) group included 83 individuals diagnosed with MCI and exhibiting Alzheimer’s disease (AD) neuropathological changes. The AD group consisted of 218 individuals with a clinical diagnosis of dementia and AD high pathology, defined as Braak stage III–VI and CERAD rated as “moderate” or “high”. An additional 94 samples categorized as “others” included individuals with a clinical diagnosis of dementia but lacking AD pathology, as well as cognitively normal individuals younger than 50 years. The Others group was included in cross-cohort calculations (Overall correlation with NPTX2 across all 575 samples) but was not included in the four-group CN-Lo/CN-Hi/MCI/AD co-expression classification.

#### PRM-MS Workflow

Human brain tissue samples selected to be representative of each of the 4 categories (CN-Lo, CN-Hi, MCI, AD) were homogenized by sonication in a lysis buffer containing 8 M urea, 50 mM triethylammonium bicarbonate (TEAB). The homogenate was centrifuged at high speed, and the resulting supernatant was collected for subsequent processing. The protein concentration of the brain lysate was determined using a bicinchoninic acid (BCA) assay. For each sample, a predetermined amount of protein was aliquoted and supplemented with synthetic isotopically-labeled peptide standards (SpikeTides™ L, JPT Peptide Technologies). These standards, featuring ¹³C- and ¹⁵N-labeled lysine and arginine residues, served as internal references for precise parallel reaction monitoring-mass spectrometry (PRM-MS) quantification. Reduction and alkylation of cysteine residues were conducted by adding tris(2-carboxyethyl) phosphine (TCEP) to a final concentration of 20 mM and incubating for 1 hour at room temperature. This was followed by alkylation with 80 mM chloroacetamide (CAA) for 30 minutes in the dark. A sequential enzymatic digestion strategy was employed for efficient protein cleavage. First, Lys-C endopeptidase (Wako Chemicals) was added at a 1:50 (w/w) enzyme-to-protein ratio and incubated for 3 hours at 37°C. The sample was then diluted with 50 mM TEAB to reduce the urea concentration to 2 M. Sequencing-grade modified trypsin (Promega) was added at a 1:50 (w/w) ratio, and digestion proceeded overnight at 37°C. The digestion was quenched by acidification with trifluoroacetic acid (TFA). The resulting peptide mixture was desalted using C18 StageTips (3M Empore™). The eluted peptides were dried in a SpeedVac vacuum concentrator (Thermo Fisher Scientific) and stored at -80°C until LC-MS/MS analysis.

Liquid chromatography-tandem mass spectrometry (LC-MS/MS) analyses were conducted based on established principles of targeted proteomics [35], with specific optimizations for the current application. An Orbitrap Fusion Lumos Tribrid mass spectrometer (Thermo Fisher Scientific), correlated to an Ultimate 3000 RSLCnano liquid chromatography system (Thermo Fisher Scientific), was used for all experiments. For chromatographic separation, tryptic peptides were first loaded onto a C18 trap column (Acclaim PepMap100, 100 μm × 2 cm, 5 μm; Thermo Fisher Scientific) at a flow rate of 5 μL/min. Analytical separation was then achieved on an EASY-Spray analytical column (50 cm × 75 μm, 2 μm C18 particles; Thermo Fisher Scientific) using a linear gradient from 6% to 28% solvent B (0.1% formic acid in 95% acetonitrile) over 55 minutes at a constant flow rate of 250 nL/min. The ion source was operated at a spray voltage of 2.0 kV. Data acquisition was performed in a targeted MS2 mode. A full MS1 scan (*m/z* 300–1600) was acquired in the Orbitrap at a resolution of 120,000 (at *m/z* 200) for profiling. Subsequently, precursor ions corresponding to the target peptides and their isotopically labeled standards were selectively fragmented via higher-energy collisional dissociation (HCD), and the resulting MS2 spectra were recorded in the Orbitrap at a resolution of 30,000. The automatic gain control (AGC) targets were set to 500,000 ions for MS1 and 100,000 ions for MS2. All raw data were processed and quantitatively analyzed using the Skyline software environment [36]. Peak areas from the extracted ion chromatograms of the target fragment ions were used to calculate the relative abundance of each endogenous peptide against its corresponding heavy internal standard for precise quantification.

To derive protein-level abundance values from peptide-level abundance values, a multi-step normalization and aggregation strategy was implemented. Initially, the raw peak area for each individual peptide was normalized to its median value across the entire sample set. Subsequently, these normalized values underwent a log2 transformation. Finally, for every target protein, the transformed values from all associated peptides were averaged to produce a single, consolidated protein abundance estimate.

### Data Analysis and Figure Generation

#### Data used for downstream analysis

Downstream analyses used a log2-transformed gene-level expression matrix (38,613 genes × 575 samples) and harmonized sample metadata (PC1, PC2, RIN, PMI, age, sex, and the NIA-AA CERAD C score and Braak B score, each on a 0–3 scale). Pathology-based grouping followed a *DoubleHi* scheme applied to cognitively normal controls: *CN-Hi* (both B score ≥ 2 and C score ≥ 2; n = 77), *CN-Lo* (the remaining controls, i.e. below 2 on either axis; n = 103), *MCI* (n = 83), *AD* (n = 218), and *Others* (n = 94). Targeted proteomic abundances for 20 targeted proteins (PRM-MS) were available for a 135-sample subset (CN-Lo n = 36, CN-Hi n = 21, MCI n = 20, AD n = 38, Others n = 20). The Others group was included in cross-cohort calculations (Overall correlation with NPTX2) but excluded from the four-group CN-Lo/CN-Hi/MCI/AD co-expression classification.

#### Targeted proteomics landscape (Figure 1)

NPTX2 PRM-MS protein and NPTX2 mRNA abundance were compared across groups with two-sided Wilcoxon rank-sum tests and BH correction (ggpubr::stat_compare_means), contrasting clinical diagnosis (Control, MCI, AD) in the left panel and neuropathology groups against a young-control reference (CN-Lo, CN-Hi, YoungCon) in the right panel; donors older than 95 years are marked with red points (Fig. 1b). For each of the 20 PRM-MS targets, Pearson correlations with NPTX2 PRM-MS protein were computed within CN-Lo, CN-Hi, MCI, and AD and rendered as a four-column correlation heatmap (Fig. 1c). In parallel, each protein’s abundance — and, in the PRM-MS samples, the corresponding gene’s mRNA — was min–max normalized to [0, 1] and summarized as group means in matched four-column expression heatmaps (Fig. 1d). Rows of all three heatmaps were ordered and split by a manually curated co-expression class. Five pairwise between-group comparisons (CN-Hi vs CN-Lo, CN-Hi vs MCI, CN-Hi vs AD, CN-Lo vs MCI, MCI vs AD) were computed using Fisher z tests on the correlations and Wilcoxon rank-sum tests on the scaled expression values, all BH-adjusted across the 20 targets.

**Figure 1.**
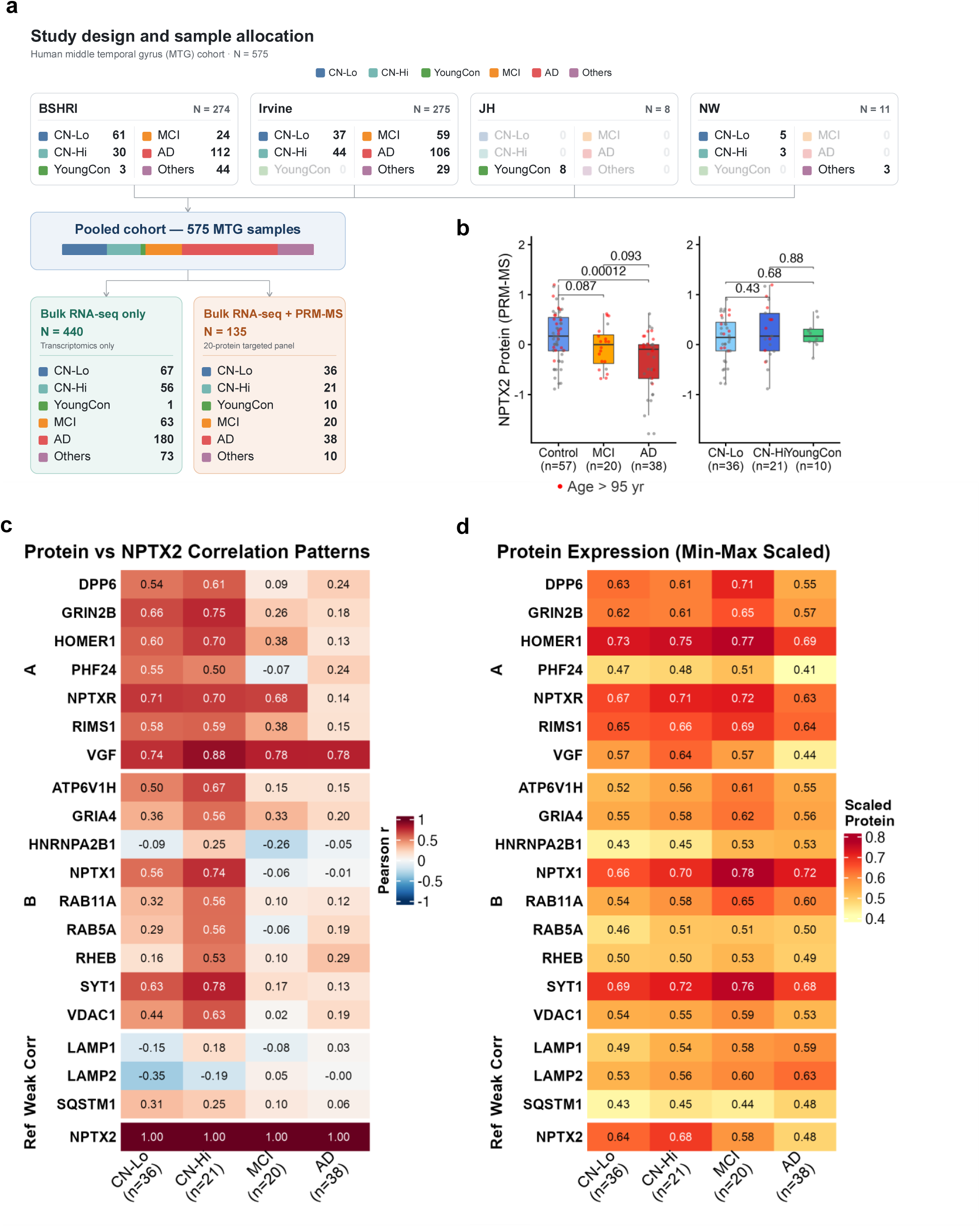
The proteomic context of the NPTX2 protein homeostasis. **a**, Study design **b**, NPTX2 protein abundance (PRM-MS) across groups, shown as box plots (center line, median; box, interquartile range (IQR); whiskers, 1.5× IQR; points, individual samples). Left, clinical diagnosis: Control (n = 57), MCI (n = 20), AD (n = 38). Right, neuropathology: CN-Lo (n = 36), CN-Hi (n = 21), YoungCon (n = 10); n denotes samples. samples older than 95 years are marked in red. Between-group comparisons were assessed by two-sided Wilcoxon rank-sum test with Benjamini–Hochberg (BH) correction across the comparisons within each panel; BH-adjusted p-values are shown on the plot. **c**, Pearson correlation of each of the 20 targeted proteins with NPTX2 protein (PRM-MS), computed within each group and displayed as a four-column heatmap of the correlation coefficient r (two-sided Pearson correlation; CN-Lo, n = 36; CN-Hi, n = 21; MCI, n = 20; AD, n = 38; n denotes samples). Rows are ordered and split by a manually curated pattern. **d**, Min–max-scaled (0–1) abundance of the 20-protein panel summarized as group means in a matched four-column heatmap, with rows in the same pattern order as **c** (CN-Lo, n = 36; CN-Hi, n = 21; MCI, n = 20; AD, n = 38; n denotes samples). CN-Hi and CN-Lo, cognitively normal controls with high and low Alzheimer’s disease neuropathology, respectively; PRM-MS, parallel reaction monitoring mass spectrometry.

#### NPTX2 protein-anchored transcriptome classification (Figure 2)

Five co-expression class schematics were generated to illustrate the allowed correlation values across CN-Lo, CN-Hi, and AD/MCI for each pattern: CN-Hi-preserved (strong in both CN-Lo and CN-Hi, weak in AD), CN-Hi-recruited (weak in CN-Lo, strong in CN-Hi, weak in AD), CN-Hi-suppressed (strong in CN-Lo, weak in CN-Hi, strong in AD), CN-Hi-reversed (strong in CN-Lo, strong with opposite sign in CN-Hi, AD either weakens or re-flips), and Pathology-disrupted (strong in CN-Lo, weak in CN-Hi and AD). Strong and weak thresholds were t_strong = 0.40 and t_weak = 0.25 (full rules in Table 1; Fig. 2a). Transcriptome-wide Pearson correlations between every gene and NPTX2 PRM-MS protein were computed within the 135-sample subset and rendered as kernel density plots segmented by correlation strength (|r| ≥ 0.5, 0.3–0.5, < 0.3) with per-group densities overlaid, accompanied beneath by a heatmap of pairwise Kolmogorov–Smirnov statistics (KS D) between groups (Fig. 2b). Stage-specific correlations of classified genes were cross-plotted as R-to-R scatterplots for two pairwise comparisons (CN-Hi vs CN-Lo and CN-Hi vs AD), with non-classified genes shown as a grey background and heatmap genes labeled (Fig. 2c, d). Classified genes were visualized as a tri-state heatmap showing, per pattern, the top five positively and top five negatively correlated genes ranked by the NPTX2 Pearson r driving the pattern assignment (Fig. 2e); red-labeled genes were co-expression-matched (same pattern and sign recovered when MCI replaced AD). Over-representation analysis used clusterProfiler::compareCluster(enrichGO, ontology = "CC", minGSSize = 15, maxGSSize = 500, BH-adjusted); pathway labels were colored red when ≥ 50% of a pathway’s hitting genes belonged to the co-expression-matched set (Fig. 2f).

**Figure 2.**
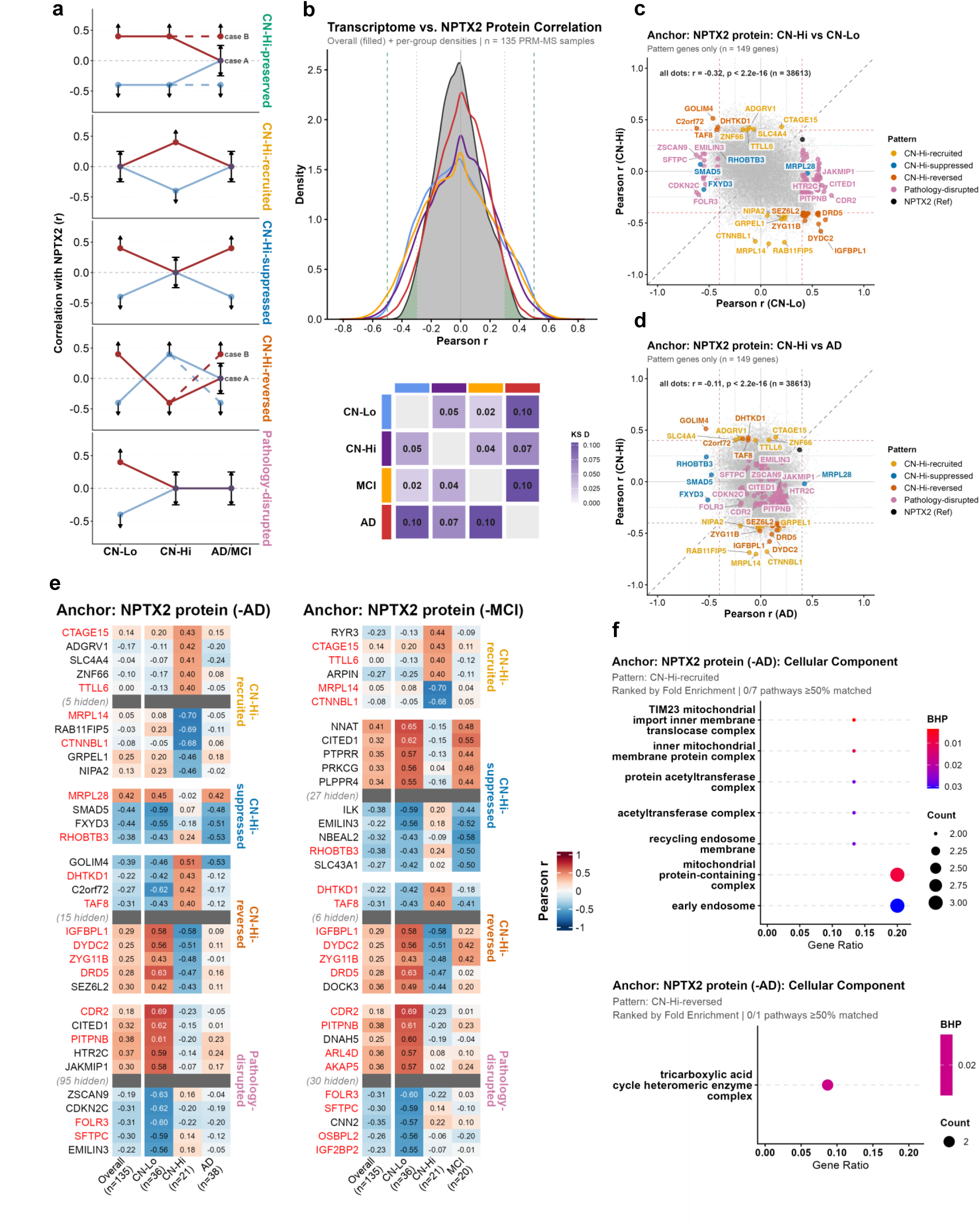
Transcriptomic Classification of NPTX2 Protein Correlated Transcripts. **a**, Schematics of the five correlation patterns defined relative to NPTX2, illustrating the permitted coefficients across CN-Lo, CN-Hi and AD/MCI (CN-Hi-preserved, CN-Hi-recruited, CN-Hi-suppressed, CN-Hi-reversed and Pathology-disrupted). Strong and weak coefficient thresholds were 0.40 and 0.25, respectively; full assignment rules are in Supplementary Table 1. No statistical analysis. **b**, Distributions of genome-wide per-gene Pearson correlation with NPTX2 protein (PRM-MS), computed within each group and shown as kernel density curves, with each gene contributing one coefficient per group (correlations computed in CN-Lo, n = 36; CN-Hi, n = 21; MCI, n = 20; AD, n = 38 samples). The lower heatmap shows the pairwise two-sided two-sample Kolmogorov–Smirnov (KS) D statistic between group distributions (n = 38,613 genes per distribution); bold entries denote Benjamini–Hochberg (BH)-adjusted p < 0.05. BH-adjusted KS p-values: CN-Lo vs CN-Hi, p = 2.3 × 10⁻³⁹; CN-Lo vs MCI, p = 9.3 × 10⁻¹⁰; CN-Lo vs AD, p = 2.3 × 10⁻¹⁶⁷; CN-Hi vs MCI, p = 3.4 × 10⁻³³; CN-Hi vs AD, p = 1.2 × 10⁻⁷⁴; MCI vs AD, p = 1.4 × 10⁻¹⁸³ **c**, Each point is a gene, positioned by its Pearson correlation with NPTX2 protein computed within CN-Lo samples (x axis, n = 36) and within CN-Hi samples (y axis, n = 21); points on the dashed diagonal have equal association in the two groups. Pattern-classified genes are colored by pattern; unclassified genes are grey. Guide lines mark coefficients of ±0.40 and ±0.25. As a measure of overall concordance between the groups, the Pearson correlation across all genes between their CN-Lo and CN-Hi coefficients — with its p-value and the number of genes — is annotated on the plot. **d**, As in **c**, for CN-Hi (y axis) versus AD (x axis) (CN-Hi, n = 21; AD, n = 38 samples). **e**, Pattern-classification heatmap (NPTX2-protein-anchored) showing, per pattern, the top five positively and top five negatively correlated genes ranked by the NPTX2 Pearson coefficient driving the assignment. Columns show the correlation coefficient r in Overall (n = 135), CN-Lo (n = 36), CN-Hi (n = 21) and AD (n = 38) samples (two-sided Pearson correlation); the disease column shows AD, with MCI (n = 20) substituted in the parallel classification. Gene labels in red are pattern-matched (same pattern and sign recovered when MCI replaced AD). **f**, Over-representation analysis (Gene Ontology Cellular Component, GO:CC) of the protein-anchored pattern gene sets, tested by one-sided hypergeometric test (clusterProfiler enrichGO) with BH correction. Dot position shows the gene ratio, dot size the number of genes, and dot colour the BH-adjusted p-value (BHP); pathway labels in red have ≥50% of their hit genes in the pattern-matched set. The Pathology-disrupted class and the MCI-substituted results are shown in Supplementary Fig. 5a. KS, Kolmogorov–Smirnov; ORA, over-representation analysis; GO:CC, Gene Ontology Cellular Component; BHP, Benjamini–Hochberg-adjusted p-value.

#### NPTX2 mRNA-anchored transcriptome classification (Figure 3)

Concordance between NPTX2 mRNA and PRM-MS protein was assessed by Pearson correlation overall and within CN-Lo, CN-Hi, MCI, and AD (135 samples; BH-corrected across five tests; Fig. 3a). NPTX2 mRNA abundance was compared across clinical diagnosis groups (Control, MCI, AD) and neuropathology groups (CN-Lo, CN-Hi, YoungCon) using Wilcoxon rank-sum tests with BH correction (Fig. 3b). Transcriptome-wide Pearson correlations between every gene and NPTX2 mRNA (575 samples) were rendered as kernel density plots as described for Figure 2b — segmented by correlation strength with per-group densities overlaid and an accompanying pairwise Kolmogorov–Smirnov (KS D) heatmap beneath — with genes reaching an overall |r| ≥ 0.5 labeled (Fig. 3c). Stage-specific correlations of classified genes were cross-plotted as R-to-R scatterplots for two pairwise comparisons — CN-Hi vs CN-Lo (Fig. 3d) and CN-Hi vs AD (Fig. 3e) — anchored on NPTX2 mRNA, with non-classified genes shown as a grey background and heatmap genes labeled. Classified genes were visualized as a pattern-classification heatmap (NPTX2-mRNA anchored) showing, per pattern, the top five positively and top five negatively correlated genes ranked by the NPTX2 Pearson r driving the pattern assignment; the third group column shows AD (with MCI substituted in the parallel validation), and red-labeled genes were trajectory-matched (Fig. 3f). Over-representation analysis of the mRNA-anchored pattern gene sets used clusterProfiler::compareCluster(enrichGO, ontology = "CC", minGSSize = 15, maxGSSize = 500, BH-adjusted), with pathway labels colored red when ≥ 50% of a pathway’s hitting genes belonged to the trajectory-matched set (Fig. 3g); additional class-specific ORA results were placed in Supplementary Fig. 5b–c.

**Figure 3.**
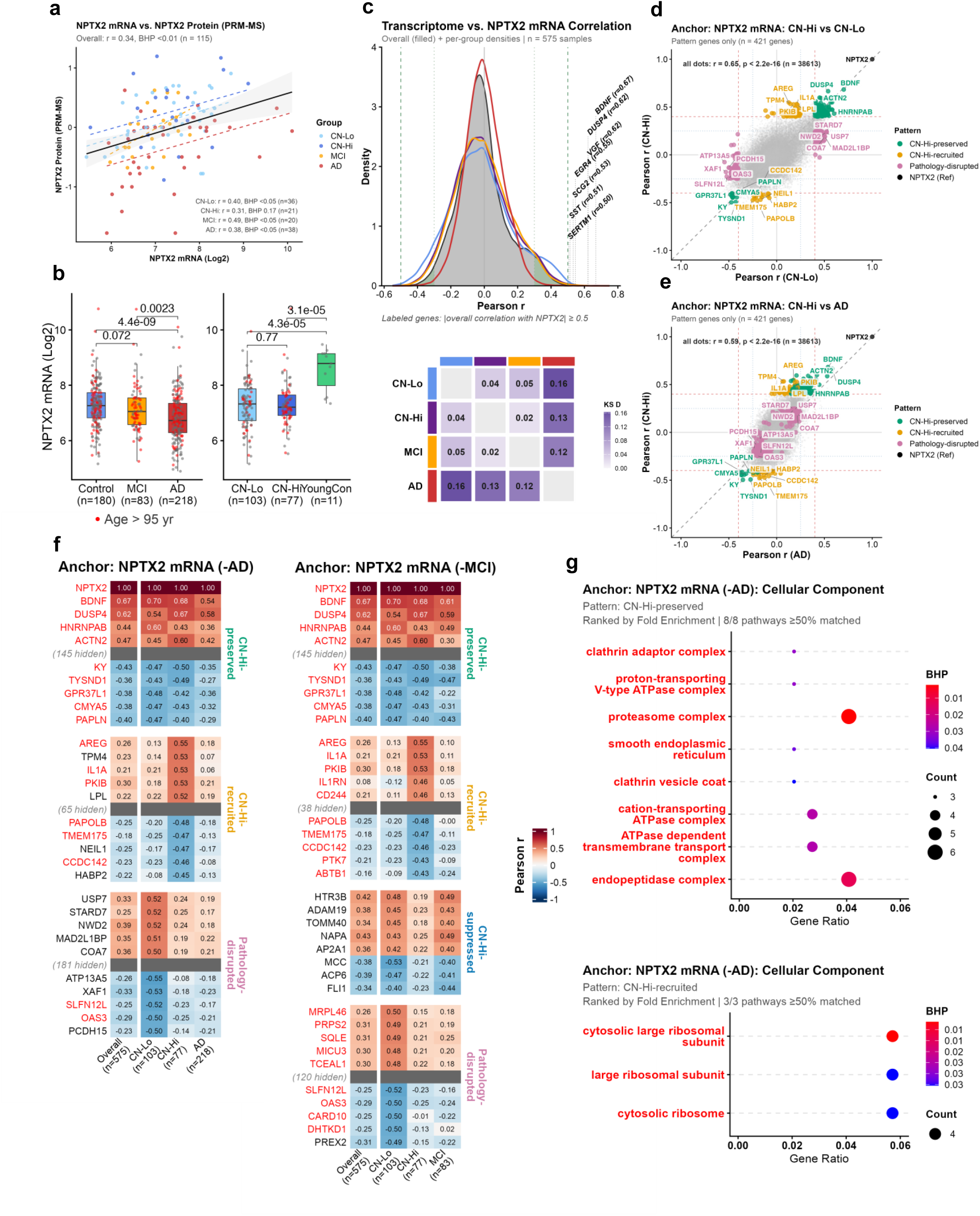
Transcriptomic Classification of NPTX2 mRNA Correlated Transcripts. **a**, NPTX2 protein abundance (PRM-MS, y axis) versus NPTX2 mRNA (log₂, x axis) in the 135-sample PRM-MS subset, colored by group; dashed lines, per-group linear fits; solid black line, overall fit (95% confidence band shaded). Association was assessed by two-sided Pearson correlation, Benjamini–Hochberg (BH)-corrected across the five tests (overall plus four groups). The overall coefficient (n = 135 samples) and the per-group coefficients (CN-Lo, n = 36; CN-Hi, n = 21; MCI, n = 20; AD, n = 38 samples), each with its BH-adjusted p-value and n, are annotated on the plot. **b**, NPTX2 mRNA abundance across groups, shown as box plots (centre line, median; box, interquartile range (IQR); whiskers, 1.5× IQR; points, individual samples). Left, clinical diagnosis: Control (n = 180), MCI (n = 83), AD (n = 218). Right, neuropathology: CN-Lo (n = 103), CN-Hi (n = 77), YoungCon (n = 11). Between-group comparisons were assessed by two-sided Wilcoxon rank-sum test with BH correction across the comparisons within each panel; BH-adjusted p-values are shown on the plot. **c**, Distributions of genome-wide per-gene Pearson correlation with NPTX2 mRNA, computed within each group and shown as kernel density curves, with each gene contributing one coefficient per group (correlations computed in CN-Lo, n = 103; CN-Hi, n = 77; MCI, n = 83; AD, n = 218 samples); genes with an overall |r| ≥ 0.5 are labelled. The lower heatmap shows the pairwise two-sided two-sample Kolmogorov–Smirnov (KS) D statistic between group distributions (n = 38,612 genes per distribution); bold entries denote BH-adjusted p < 0.05. BH-adjusted KS p-values: CN-Lo vs CN-Hi, p = 9.8 × 10⁻²⁴; CN-Lo vs MCI, p = 1.9 × 10⁻⁴²; CN-Lo vs AD, p < 1 × 10⁻³⁰⁰; CN-Hi vs MCI, p = 4.6 × 10⁻⁶; CN-Hi vs AD, p = 5.4 × 10⁻³⁰²; MCI vs AD, p = 1.8 × 10⁻²²⁸. **d**, Each point is a gene, positioned by its Pearson correlation with NPTX2 mRNA computed within CN-Lo samples (x axis, n = 103) and within CN-Hi samples (y axis, n = 77); points on the dashed diagonal have equal association in the two groups. Pattern-classified genes are colored by pattern; unclassified genes are grey. Guide lines mark coefficients of ±0.40 and ±0.25. As a measure of overall concordance between the groups, the Pearson correlation across all genes between their CN-Lo and CN-Hi coefficients — with its p-value and the number of genes — is annotated on the plot. **e**, As in **d**, with the per-gene NPTX2-mRNA correlation computed within AD samples (x axis, n = 218) and within CN-Hi samples (y axis, n = 77). **f**, Pattern-classification heatmap (NPTX2-mRNA-anchored) showing, per pattern, the top five positively and top five negatively correlated genes ranked by the NPTX2 Pearson coefficient driving the assignment. Columns show the correlation coefficient r in Overall (n = 575), CN-Lo (n = 103), CN-Hi (n = 77) and AD (n = 218) samples (two-sided Pearson correlation); the disease column shows AD, with MCI (n = 83) substituted in the parallel classification. Gene labels in red are pattern-matched (same pattern and sign recovered when MCI replaced AD). **g**, Over-representation analysis (Gene Ontology Cellular Component, GO:CC) of the mRNA-anchored pattern gene sets, tested by one-sided hypergeometric test (clusterProfiler enrichGO) with BH correction. Dot position shows the gene ratio, dot size the number of genes, and dot colour the BH-adjusted p-value (BHP); pathway labels in red have ≥50% of their hit genes in the pattern-matched set. The Pathology-disrupted class and the MCI-substituted results are shown in Supplementary Fig. 5b, c.

#### Transcriptome-wide NPTX2 circuit classification (Figures 2–4)

Pearson correlations between every gene and NPTX2 were computed within CN-Lo, CN-Hi, AD, and MCI at both the RNA level (gene mRNA vs NPTX2 mRNA, 575 samples) and the cross-omic level (gene mRNA vs NPTX2 PRM-MS protein, 135-sample subset). For the master gene list, the maximum absolute r across six group-by-modality combinations (CN-Lo, CN-Hi, and AD × RNA and Protein — excluding MCI from the selection pool) was retained with its sample size. Two-sided p-values (t distribution, n − 2 df) were BH-adjusted transcriptome-wide. Genes were retained if mean log2 expression ≥ 1.0, maximum |r| > 0.40, and BH-adjusted p < 0.01, after filtering to protein-coding transcripts via biomaRt (16,434 valid symbols) [39].

**Figure 4.**
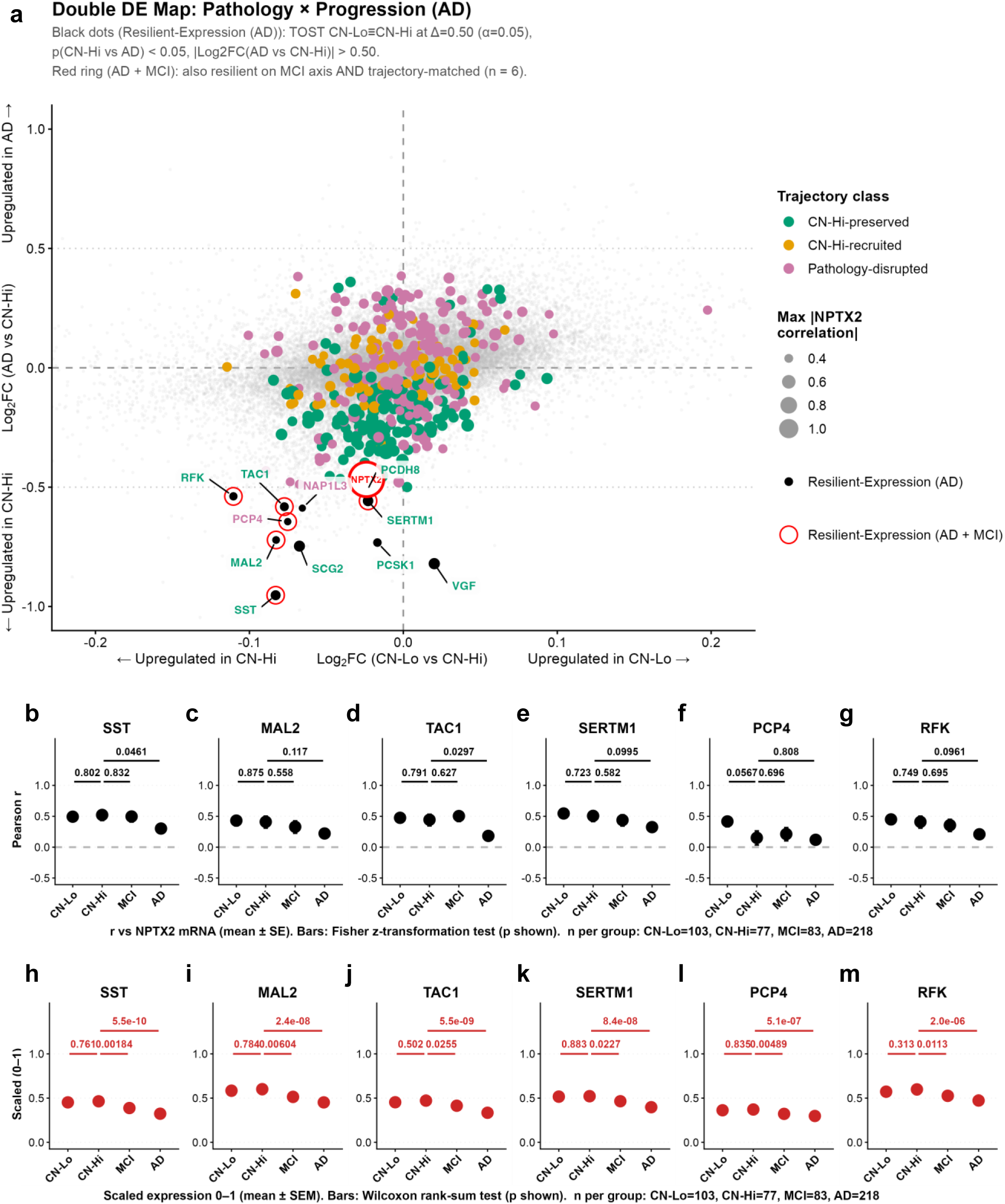
NPTX2-Associated Transcripts with Preserved Expression in Asymptomatic Groups. **a**, Double differential-expression map; each point is a gene. The x axis is the log₂ fold change between CN-Lo and CN-Hi (equivalence axis) and the y axis the log₂ fold change between AD and CN-Hi (progression axis). Grey points, all remaining genes (background); colored points, pattern-classified genes (colored by pattern, point size proportional to the maximum absolute NPTX2 correlation). Filled black points are resilient-expression genes — equivalent between CN-Lo and CN-Hi by two one-sided tests (TOST; equivalence bound Δ = 0.50 log₂ units, α = 0.05) and significantly different between CN-Hi and AD by two-sided Welch’s t-test (p < 0.05, |log₂ FC| > 0.50). Red rings mark the subset that additionally met resilient-expression criteria against the MCI reference and was pattern-matched (n = 6 genes); the red point marks NPTX2. Tests used n = 103 (CN-Lo), 77 (CN-Hi), 218 (AD) and 83 (MCI) samples. Dotted guides mark Δ = ±0.50 (x) and |log₂ FC| = 0.50 (y). Panels **b**–**g** (correlation) and **h**–**m** (expression) show the six resilient-expression genes — those whose NPTX2 association and expression are preserved between CN-Lo and CN-Hi but altered in MCI and AD — with the gene indicated above each panel. **b**–**g**, For each gene, the Pearson correlation between its mRNA and NPTX2 mRNA computed within each group, shown as one point per group (CN-Lo, n = 103; CN-Hi, n = 77; MCI, n = 83; AD, n = 218 samples); error bars are ±1 standard error of the Fisher z-transformed coefficient, back-transformed to the correlation scale. Pairwise comparisons (CN-Lo vs CN-Hi, CN-Hi vs MCI, CN-Hi vs AD) were assessed by a two-sided Fisher z-transformation test; p-values are shown on the plot. **h**–**m**, For the same six genes, min–max-scaled (0–1) mRNA expression within each group, shown as box plots (centre line, median; box, interquartile range; whiskers, 1.5× IQR) with individual samples overlaid (CN-Lo, n = 103; CN-Hi, n = 77; MCI, n = 83; AD, n = 218). Pairwise comparisons (CN-Lo vs CN-Hi, CN-Hi vs MCI, CN-Hi vs AD) were assessed by two-sided Wilcoxon rank-sum test; p-values are shown on the plot. TOST, two one-sided tests.

Filtered genes were assigned to one of five co-expression patterns — CN-Hi-preserved, CN-Hi-recruited, CN-Hi-suppressed, CN-Hi-reversed, and Pathology-disrupted — based on the direction and magnitude of their stage-specific correlations using thresholds t_strong = 0.40 and t_weak = 0.25 (Supplementary Table 1). The classification was run independently using NPTX2 mRNA and NPTX2 PRM-MS protein as anchors, and in parallel using AD and MCI as the third disease group, yielding four classifications (RNA/AD, RNA/MCI, Protein/AD, Protein/MCI).

Genes whose pattern and sign were concordant between the AD-based and MCI-based classifications were flagged as co-expression-matched.

#### Resilient-expression analysis (Figure 4)

To isolate genes whose expression shifts specifically between the symptomatic groups (CN-Hi vs AD) while remaining equivalent between the pre-symptomatic pair (CN-Lo vs CN-Hi), we built a double differential expression map with log2(CN-Lo − CN-Hi) on the x axis and log2(AD − CN-Hi) on the y axis (Fig. 4a). Equivalence on the x-axis was established by two one-sided tests (TOST) with an equivalence bound Δ = 0.50 log2 units at α = 0.05, requiring that the 90% CI of the CN-Lo − CN-Hi mean difference fall entirely within ±Δ. A gene was flagged as having resilient expression on the AD axis when it passed TOST equivalence and showed a significant Welch t-test on the y-axis (p < 0.05, |log2 FC| > 0.50). An equivalent analysis was performed with MCI replacing AD on the y-axis (no |FC| filter on MCI). Genes meeting resilient-expression criteria on both axes and belonging to the co-expression-matched set from the parallel AD/MCI classification were highlighted with a red open ring and labeled. For the six resilient-expression candidates that met both criteria and belonged to the co-expression-matched set under both disease references — genes whose NPTX2 association is preserved in CN-Hi but disrupted in MCI and AD — per-group statistics versus NPTX2 mRNA were plotted across CN-Lo, CN-Hi, MCI, and AD as two panel rows: Pearson correlations, each shown as a point per group with error bars spanning ±1 SE on the Fisher z scale back-transformed to the −1 to 1 axis (Fig. 4b–g), and min–max scaled expression (0–1), shown as boxplots (median and IQR) with individual samples overlaid (Fig. 4h–m). Groups were not connected by lines, and the CN-Lo vs CN-Hi, CN-Hi vs MCI, and CN-Hi vs AD comparisons were annotated using Fisher z difference tests for the correlations and Wilcoxon rank-sum tests for the expression values.

### Supplementary analyses

Differential expression between CN-Hi and CN-Lo, and between AD and CN-Hi, was assessed for the 20 PRM-MS proteins using per-protein Wilcoxon rank-sum tests (BH-adjusted, FDR < 0.05, |log2 FC| > 0.20) and presented as volcano plots (Supplementary Fig. 1a, 1b). Pairwise Pearson correlations between the 20 PRM-MS proteins and clinical metadata (age, PMI, CERAD, Braak) were computed in the 135-sample PRM-MS subset and presented as a heatmap of the metadata columns only (Supplementary Fig. 1c). The same contrasts were tested across ∼16,000 protein-coding transcripts using limma (eBayes, trend = FALSE, robust = TRUE; BH-adjusted, FDR < 0.01, |log2 FC| > 0.50), with NPTX2 labeled in the volcano plots (Supplementary Fig. 4a, 4b). The precomputed PC1–PC2 scatter was used to visualize PRM-MS subset membership, shown as hollow red rings over all 575 samples (Supplementary Fig. 2a).

The biological identity of PC1 and PC2 was characterized by fgsea against GO:CC gene sets (minSize = 50, maxSize = 1000; top 10 positively and negatively enriched terms per component), presented as compact bar plots (Supplementary Fig. 2b, 2c). Pairwise Pearson correlations between the 20 targeted gene mRNAs (plus PC1 and PC2) and clinical metadata (RIN, age, PMI, CERAD, Braak) were computed across all 575 samples and presented as a heatmap of the metadata columns only (Supplementary Fig. 4c). To evaluate potential sex confounding, distributions of transcriptome-wide gene–NPTX2 mRNA Pearson correlations were compared between Female and Male donors within each group using two-sample Kolmogorov–Smirnov tests (Supplementary Fig. 3a–d). Sex balance across clinical covariates (age, PMI, CERAD, Braak, RIN) and gene-level |Pearson r| was assessed by Wilcoxon rank-sum tests with BH correction within each panel (Supplementary Fig. 3e–j). Over-representation analyses (GO:CC) for individual co-expression classes not shown in the main-figure ORA panels are provided separately: the Pathology-disrupted class from the NPTX2-protein-anchored classification (Supplementary Fig. 5a), and the CN-Hi-preserved and CN-Hi-recruited classes from the NPTX2-mRNA-anchored classification (Supplementary Fig. 5b, 5c).

### Statistical software and reproducibility

All analyses were performed in R 4.4.2 (2024-10-31) on macOS (aarch64-apple-darwin20). Correlations were computed with base R *cor* and *cor.test* (method = "pearson", use = "pairwise.complete.obs"). Group comparisons used *wilcox.test* and *t.test*; equivalence testing used two one-sided *t*-tests (TOST). Multiple-testing correction throughout was Benjamini–Hochberg via *p.adjust*. Gene set enrichment used *fgsea*; over-representation analysis used *clusterProfiler* with *org.Hs.eg.db*. Protein-coding annotation was obtained from Ensembl via *biomaRt* (hsapiens_gene_ensembl). Differential expression for RNA used *limma* (eBayes with robust moderation). Heatmaps were rendered with *ComplexHeatmap* and *circlize*. Data manipulation used *dplyr*, *tidyr*, *tibble*, *readr*, *readxl*, *writexl*, and *stringr*. Visualizations were produced with *ggplot2*, *ggpubr*, *ggrepel*, *scales*, *RColorBrewer*, *cowplot*, *gridExtra*, and *svglite*. Figures were rendered in Arial and exported as SVG.

## Results

### Cohort sourcing and processing

Analysis was performed on postmortem middle temporal gyrus (MTG) tissue collected from four independent cohorts (Fig. 1A). The Johns Hopkins cohort contributed young controls, the Banner cohort contributed young controls, cognitively normal controls that included LoPath and HiPath, MCI, AD, and other pathology cases, the Irvine cohort contributed cognitively normal controls, MCI, AD, and other pathology cases, and the SuperAging Research Initiative contributed cognitively average controls.

After quality control, 135 representative samples spanning the diagnostic groups were selected for PRM-MS proteomic analysis and bulk RNA-seq. The RNA-seq cohort was subsequently expanded with 440 additional samples, yielding a total of 575 MTG transcriptomes.

#### NPTX2 Protein Correlation with Targeted Proteomic Panel

Control samples were subdivided into low-pathology (CN-Lo) and high-pathology (CN-Hi) groups according to CERAD neuritic-plaque and Braak neurofibrillary-tangle criteria (NIA-AA C and B scores, each on a 0–3 scale). Controls were classified as CN-Hi when both the B score and the C score were ≥ 2, and as CN-Lo otherwise. PRM-MS analysis demonstrated that NPTX2 protein abundance was stable across all control groups and reduced in MCI and AD groups (Fig. 1b).

To define the molecular context associated with NPTX2 protein abundance, we quantified 19 additional proteins representing: (i) proteins linked to NPTX2 through physical interaction or prior reported association (NPTX1, NPTXR, GRIA4, VGF, SYT1, HNRNPA2B1, ATP6V1H, GRIN2B, RIMS1 [2, 3, 14, 16, 37–46]); (ii) proteins associated with cognitive trajectory (VDAC1, DPP6, PHF24/KIAA1045, HOMER1[48–52]); and (iii) proteins associated with autophagy–lysosomal pathways (SQSTM1/P62, LAMP1, LAMP2, RAB11A, RAB5A), together with RHEB, a regulator of lysosome-associated mTORC1 signaling and cellular metabolism (Fig. 1c).

There is a clear loss of proteome correlation structure centered on NPTX2 in MCI and AD. This NPTX2 network is especially tightly coordinated specifically in CN-Hi controls (Fig. 1C). In addition to validating this networks’ link to in cognitive decline, this also suggests that CN-Hi and CN-Lo may navigate distinct molecular trajectories to preserve cognition. Consistent with this possibility, for LAMP1 and HNRNPA2B1 the direction of correlation with NPTX2 is opposite within CN-Hi and within CN-Lo subjects (Fig. 1c). This may indicate a coupling of autophagy-lysosomal pathways and synaptic gene expression specifically in CN-Hi.

Several proteins showed stronger correlation with NPTX2 protein specifically in CN-Hi. This, comprised synaptic and proteostasis components NPTX1, GRIA4, SYT1, RAB5A, RAB11A, ATP6V1H, RHEB, and VDAC1 (class B; Fig. 1c). By contrast, another set (class A; Fig. 1c) maintained strong correlation with NPTX2 across both control groups before weakening in MCI and then further in AD. This set was enriched for synaptic and inhibitory systems, comprising the pentraxin–AMPA–NMDA signaling complex (NPTXR, GRIN2B, HOMER1), presynaptic release machinery (RIMS1), dendritic excitability regulators (DPP6), and inhibitory signaling proteins (PHF24). Interestingly, the neurotrophin-responsive peptide precursor VGF exhibited consistent strong correlation with NPTX2 even in AD, but with slight elevation in CN-Hi.

In contrast to this loss of correlation structure, differences in protein abundance were not clear across diagnostic groups (Fig. 1d). NPTX2 itself was an exception, with reduced levels in AD (Fig. 1b, d). VGF was the only protein concordantly elevated in CN-Hi in both abundance and NPTX2 correlation.

#### Transcriptome-Wide Co-Expression Anchored on NPTX2 Protein

In the transcriptome-wide analysis we applied the same co-expression framework using correlations between transcript abundances and NPTX2 protein level. Five co-expression classes were defined: CN-Hi-preserved, CN-Hi-recruited, CN-Hi-suppressed, CN-Hi-reversed, and pathology-disrupted (Fig. 2a). Expression of genes in distinct NPTX2 protein co-expression classes can be freely explored in our bulk RNA-seq data and other public AD transcriptomic datasets at **NeMO Analytics**.

Transcriptome-wide density plots showed reduced transcriptomic correlations with NPTX2 protein in AD relative to CN-Lo, CN-Hi, and MCI distributions (Fig. 2b). This indicates that NPTX2-associated transcriptomic coordination is not globally reduced in CN-Hi or MCI. Nevertheless, pairwise R–R scatterplots across the CN-Lo-to-CN-Hi and CN-Hi-to-AD comparisons resolved discrete CN-Hi-recruited, CN-Hi-reversed, and CN-Hi-suppressed transcript subsets (Fig. 2c–d). Across the transcriptome, per-transcript NPTX2 protein correlations showed low cross-group concordance: they were inversely related between CN-Hi and CN-Lo (r = −0.32, p < 2.2 × 10⁻¹⁶) and near-orthogonal between CN-Hi and AD (r = −0.11, p < 2.2 × 10⁻¹⁶), indicating that the NPTX2 co-expression structure in CN-Hi is distinct from that in both CN-Lo and AD. This distinct structure shows that NPTX2-associated molecular rewiring was readily detectable in CN-Hi individuals.

Transcriptome-wide analysis identified 15 CN-Hi-recruited, 4 CN-Hi-suppressed, and 24 CN-Hi-reversed transcripts (Fig. 2e). Over-representation analysis of the CN-Hi-recruited class returned a mitochondrial inner-membrane and import signature (Fig. 2f), but this enrichment was not robust: its driving transcripts were largely not recovered when MCI was substituted for AD as the disease reference group — that is, not assigned the same CN-Hi co-expression class under both references. Four of the 15 recruited transcripts were recovered under both AD and MCI references, spanning mitochondrial translation (MRPL14), ER-cargo exit (CTAGE15), microtubule glutamylation (TTLL6), and pre-mRNA splicing (CTNNBL1). Of the four suppressed transcripts, two were recovered under both references — MRPL28 (mitochondrial translation) and RHOBTB3 (Golgi-to-ER transport). The CN-Hi-reversed class was the largest, but its sole enriched term, the tricarboxylic-acid-cycle enzyme complex (Fig. 2f), was likewise not robust to the reference substitution between MCI and AD; of its 24 transcripts, 12 were recovered under both references, spanning energy metabolism (GOT1, DHTKD1), microtubule and cytoskeletal regulation (MAP1B, NUAK1), proteostasis (PSMA5, ZYG11B), ER–Golgi trafficking (TRAPPC3), dopaminergic neuromodulation (DRD5), cell adhesion (CDH13), growth-factor signaling (IGFBPL1), transcription (TAF8), and a poorly characterized transcript (DYDC2). Because MCI is closer than AD to CN-Hi in pathology severity (Supplementary Fig. 3g, h), transcripts recovered under both the AD and MCI references retain their CN-Hi–specific co-expression independent of pathology burden, indicating that these associations are tied to the CN-Hi resilience state itself rather than to disease severity. These span a wide functional range—from mitochondrial translation and energy metabolism to cytoskeletal regulation, proteostasis, and ER–Golgi trafficking—rather than a single pathway.

A total of 105 transcripts were classified as pathology-disrupted, only 34 of which were also recovered in the MCI-based classification. Over-representation analysis identified enrichment of synaptic gene sets within this category (Supplementary Fig. 5a).

#### Transcriptome-Wide Co-Expression Anchored on NPTX2 mRNA

We next applied an NPTX2 mRNA-anchored co-expression framework to the expanded 575-sample MTG RNA-seq cohort. Principal component analysis confirmed that the 135-sample PRM-MS subset was representative of the expanded cohort and additionally identified associations between targeted transcripts and tissue-quality variables, including postmortem interval and RIN (Supplementary Fig. 2a; Supplementary Fig. 4c). Despite the larger cohort size compared to the proteomics, a direct differential expression analysis between CN-Lo and CN-Hi identified no significant differentially expressed transcripts (Supplementary Fig. 4a-b).

Accordingly, we examined the possibility that the resilient state was more strongly characterized by altered molecular coordination with NPTX2 mRNA than by broad transcript abundance changes.

NPTX2 mRNA abundance was moderately correlated with NPTX2 PRM-MS protein abundance across the 135-sample subset (Fig. 3a). Like NPTX2 protein, NPTX2 mRNA was similar across CN-Lo and CN-Hi groups and reduced in MCI and AD. However, unlike NPTX2 protein, which remained stable across asymptomatic aging, NPTX2 mRNA abundance was significantly lower in aged controls relative to young controls (Fig. 3b).

As with correlations with NPTX2 protein (Fig. 2b), transcriptome-wide density plots showed reduced correlations with NPTX2 mRNA in AD, whereas the CN-Lo, CN-Hi, and MCI distributions remained largely overlapping (Fig. 3c). Sex-stratified density plots further showed that the AD-associated collapse in correlation strength was more pronounced in females in all pathological conditions (CN-Hi, MCI, and AD), while in CN-Lo, correlations were slightly higher in females than males (Supplementary Fig. 3). Several activity-dependent and synaptic transcripts, including BDNF, VGF, SCG2, SST, SERTM1, DUSP4, and EGR4, showed strong overall correlations with NPTX2 mRNA and formed a CN-Hi-preserved co-expression class that was not observed in the protein-anchored analysis (Fig. 3c). Several of these transcripts are well established in neurotrophic and activity-dependent neuropeptide signaling—BDNF [53], and SCG2, which is induced together with VGF in the parvalbumin-interneuron in response to activity [54]. Pairwise R–R scatterplots across the CN-Lo-to-CN-Hi and CN-Hi-to-AD transitions further resolved the resilience-associated co-expression structure. Per-transcript NPTX2 correlations in CN-Hi were strongly positively correlated with those in both CN-Lo (r = 0.65, p < 2.2 × 10⁻¹⁶) and AD (r = 0.59, p < 2.2 × 10⁻¹⁶), indicating that the overall NPTX2 co-expression architecture is broadly preserved across groups, with CN-Hi remaining somewhat more concordant with CN-Lo than with AD (Fig. 3d–e). Expression of genes in distinct NPTX2 mRNA co-expression classes can be freely explored in our bulk RNA-seq data and other public AD transcriptomic datasets at **NeMO Analytics**.

Transcriptome-wide analysis identified 154 CN-Hi-preserved, 75 CN-Hi-recruited, and 192 pathology-disrupted transcripts (Fig. 3f), with the preserved class comprising 141 positively correlated transcripts (excluding NPTX2) and 13 negatively correlated transcripts. Over-representation analysis of the CN-Hi-preserved class identified enrichment of proteasome/peptidase, clathrin-coated vesicle and adaptor, V-type ATPase, ER-lumen, transport-vesicle, and focal-adhesion/cell-substrate-junction pathways (Fig. 3g); the ER-lumen and transport-vesicle terms reflect a coherent activity-induced secretory program of neurotrophin and neuropeptide cargo (BDNF, VGF, and SCG2, together with PCSK1 and ADCYAP1). When the 141 positively correlated preserved partners were grouped by biological function, proteostasis-related machinery (spanning protein synthesis, folding, trafficking, lysosomal function, and degradation) formed the largest category (68 of 141 partners), while the single largest discrete program was an activity-dependent immediate-early transcription-factor and feedback-regulator network (17 of 141; EGR family members, FOSB, NPAS4, DUSP phosphatases, and RGS2), all remaining tightly associated with NPTX2 across asymptomatic states. The CN-Hi-recruited class was enriched for cytosolic ribosomal-subunit pathways (Fig. 3g) and additionally included a prominent inflammatory, neuroprotective, and immune-regulatory component, with cytokines, chemokines, and immune regulators (IL1A, CCL3, CCL4, CXCL8, IL1RN, AREG, LIF, CH25H, and GPR183) becoming selectively coordinated with NPTX2 in CN-Hi individuals.

Parallel classification using MCI in place of AD recovered 60% of pathology-disrupted transcripts and 64% of CN-Hi-recruited transcripts (Fig. 3f). Together, the CN-Hi–specific NPTX2 mRNA co-expression structure implicates a preserved proteostasis and activity-dependent immediate-early program alongside a CN-Hi–recruited ribosomal and immune-regulatory component, spanning a broad functional range rather than a single pathway.

#### Identification of NPTX2-Associated Resilient-Expression Candidates

To capture cognition-associated programs that are disrupted in AD and MCI, but whose molecular integrity is preserved in high-pathology cognitively normal individuals, we performed an analysis to identify transcripts that remained stable in CN-Hi – in both expression and coordination with NPTX2 –but altered in symptomatic conditions.

We first compared differential expression between CN-Lo and CN-Hi against differential expression between CN-Hi and AD. Candidate transcripts were required to show unchanged expression between CN-Lo and CN-Hi by two one-sided tests (TOST; Δ = 0.5, α = 0.05; Fig. 4a X-axis), together with significant differential expression between CN-Hi and AD (Fig. 4a Y-axis). This identified 11 resilient-expression candidates (labeled genes in black in Fig. 4a). 6 of these candidates were also identified after substituting AD with MCI (genes circled in red in Fig. 4a). Among the 6 recovered candidates, 5 displayed CN-Hi-preserved correlation trajectories with NPTX2 mRNA: SST, MAL2, TAC1, SERTM1, and RFK [55–57]. The remaining one candidate, PCP4, displayed pathology-disrupted correlation trajectories (Fig. 4b–n).

## Discussion

Cognitive resilience to AD pathology is increasingly understood as a systems-level phenomenon in which cognition is preserved despite substantial accumulation of Aβ plaques and neurofibrillary tangles. Among molecular correlates of resilience, NPTX2 has emerged as one of the most reproducible single-gene markers, with stable brain and CSF levels associated with preserved cognition independent of classical AD biomarkers [9, 58]. These observations suggest that NPTX2-associated biology might serve as a point of reference to identify broader mechanisms implicated in cognitive resilience despite the presence of AD neuropathology.

To address this question, we used a NPTX2-centered co-expression framework rather than conventional differential-expression analysis. Differential expression between CN-Lo and CN-Hi groups was minimal, indicating that resilience is not primarily underpinned by large shifts in transcript abundance (Supplementary Fig. 1, 4). This framework resolved two main resilience-associated co-expression classes: the CN-Hi-preserved class identified molecular relationships maintained across low- high-pathology states in cognitively normal controls, whereas the CN-Hi-recruited class captured relationships selectively engaged in cognitively resilient individuals with high pathology burden.

In the targeted protein analysis, the CN-Hi-preserved class (class A, Fig. 1c) comprised excitatory and inhibitory synaptic relationships that remained coordinated with NPTX2 across both low- and high-pathology cognitively normal individuals, weakening only with cognitive impairment. With the exception of VGF, these preserved protein–NPTX2 relationships were not reproduced by the same genes at the transcript level; their synaptic functional architecture was nonetheless recapitulated in the transcript analysis and extended into a broader preserved program. Within this CN-Hi-preserved transcript-level program, proteostasis-related machinery formed the dominant component, alongside a prominent activity-dependent immediate-early transcriptional network (Fig. 3). Together these observations suggest that resilient brains preserve coordinated synaptic organization at the protein level and an expanded protein-homeostasis and activity-dependent transcriptional network at the transcript level, despite substantial neuropathology. In addition to NPTX2’s well-known function in maintaining PV function, this CN-Hi preserved network also extended to other inhibitory-circuit components: somatostatin (SST) remained coordinated with NPTX2 in resilient individuals. This is important as SST interneurons are among the earliest and most vulnerable neuronal populations in AD [59, 60].

In contrast, the CN-Hi-recruited class comprised relationships selectively engaged in resilient individuals with high pathology burden. In the targeted protein panel (Fig.1c), the recruited partners included synaptic-release components together with autophagy–lysosomal and mTORC1-associated proteins, each strengthening its correlation with NPTX2 specifically in high-pathology resilient individuals; this autophagy–lysosomal arm of proteostasis was thus engaged specifically in the context of high pathology, in contrast to the proteasomal machinery that remained preserved across both CN-Lo and CN-Hi. At the transcript level (Fig. 2-3), the CN-Hi recruited relationships reproduced under both disease references engaged inflammatory cytokine and chemokine signaling together with ribosomal and metabolic machinery. The co-recruitment of pro-inflammatory mediators alongside their endogenous antagonists and neuroprotective regulators suggests that resilience is not a state of immune quiescence but one of regulated neuroimmune coordination with NPTX2 [61], while the concurrent translational and metabolic engagement indicates that resilient states may actively mobilize biosynthetic and energetic capacity as pathology increases [62] .

Importantly, CN-Hi-specific remodeling extended beyond newly recruited relationships: a smaller set of associations was selectively suppressed or reversed in CN-Hi (Fig. 2f). Together, these patterns indicate that resilience is not a single process but encompasses multiple elements of NPTX2-associated reorganization including both the recruitment of new relationships and the uncoupling of existing ones.

Multi-modal convergence further elucidated the resilience framework. Across analyses, two partners converged. VGF, proposed as a biomarker and therapeutic target in AD [63, 64], was the strongest convergent signal, preserved in its correlation with NPTX2 at both the protein and transcript levels and uniquely elevated in absolute abundance in CN-Hi individuals. CTAGE15 was recruited into NPTX2 co-expression in mRNA analyses anchored to both NPTX2 protein and mRNA, thereby linking resilience to ER-export systems. Together with the distinct CN-Hi preserved, recruited, suppressed, and reversed patterns described above, these observations argue that resilience-associated relationships undergo selective remodeling rather than uniform activation.

A dominant systems-level feature of the preserved resilience network was the prominence of proteostasis, a system that has being shown to be disrupted in aging and neurodegenerative diseases [62]. Although NPTX2 is generally conceptualized as a synaptic protein, its preserved transcript-level co-expression partners belonged most heavily to pathways regulating protein synthesis, folding, trafficking, lysosomal function, and degradation (Fig. 3g), collectively exceeding any single other program even though activity-dependent transcription was the largest individual one. This breadth is biologically compelling because the same machinery required for NPTX2 biosynthesis and turnover also mediates clearance of misfolded Aβ and tau [65]: NPTX2 is a secreted glycoprotein requiring ER folding, glycosylation, vesicular trafficking, and lysosomal processing [14], placing it directly within the proteostasis systems most vulnerable during AD progression.

The present data do not resolve whether these proteostasis relationships reflect intrinsic constitutional resilience or adaptive recruitment under pathology burden [66]. Both interpretations remain plausible. Nonetheless, the prominence of proteostasis within the preserved class, together with its re-emergence as an autophagy–lysosomal arm under recruitment, argues that maintenance of protein homeostasis is central to resilient aging.

Finally, five resilient-expression candidate biomarkers (SST, TAC1, MAL2, SERTM1, and RFK) distinguished CN-Hi from both AD and MCI in abundance while remaining unchanged relative to CN-Lo (Fig. 4a). These span inhibitory and neuropeptidergic signaling, membrane trafficking, and metabolism, reinforcing the distributed nature of the resilience network.

Several limitations should be acknowledged. The study is cross-sectional and therefore cannot establish causality or temporal sequence. Correlation trajectories do not distinguish direct regulation from shared upstream control, and bulk-tissue analyses cannot resolve cell-type-specific contributions. In addition, sex may add a further layer as shown above (Supplementary Fig. 3), the loss of NPTX2-associated coordination under pathology was more pronounced in females, yet sex was not explicitly modeled in the main co-expression framework. Future studies that utilize single-cell analyses [67] or approaches that can combine longitudinal sampling or perturbation approaches will be required to determine how these resilience-associated networks are established and maintained.

In summary, resilience is organized as a layered molecular network defined by relationships with NPTX2 that are selectively preserved, recruited, or remodeled. The preserved class maintains the basal architecture of synaptic organization and proteostatic homeostasis across asymptomatic aging, whereas the recruited class engages adaptive immune, translational, and metabolic programs specifically under high pathology burden. Proteostasis adaptation recurred across these layers, dominating the preserved network as a proteasomal and biosynthetic program, making protein-homeostasis the most pervasive systems-level feature of the cognitively resilient brain.

## Conclusion

By combining targeted proteomics with transcriptome-wide analyses, we developed a co-expression-based framework to characterize molecular coordination around NPTX2 across CN-Lo, CN-Hi, MCI, and AD. This framework showed that cognitive resilience is not primarily defined by broad transcript abundance changes, but by preserved and selectively remodeled relationships with NPTX2. Structural synaptic and inhibitory-signaling proteins remained coordinated with NPTX2 across asymptomatic groups, whereas CN-Hi-specific trajectories implicated adaptive recruitment, uncoupling, and rewiring of pathways involved in trafficking, proteostasis, metabolic signaling, immune regulation, and neuromodulation. Transcriptome-wide analyses further identified resilient-expression candidates, including SST, MAL2, TAC1, SERTM1, and RFK, that remained preserved in CN-Hi but differed in MCI and AD. Together, these findings position NPTX2 as a central node within a broader resilience-associated molecular network and provide a framework for future studies of cognitive resilience in AD.

## List of abbreviations

Aβ: amyloid-β
AD: Alzheimer’s disease
AMPA: α-amino-3-hydroxy-5-methyl-4-isoxazolepropionic acid
BH: Benjamini–Hochberg (correction)
CERAD: Consortium to Establish a Registry for Alzheimer’s Disease
CI: confidence interval
CN-Hi: cognitively normal, high-pathology (control)
CN-Lo: cognitively normal, low-pathology (control)
CSF: cerebrospinal fluid
ER: endoplasmic reticulum
FC: fold change
FDR: false discovery rate
GO:CC: Gene Ontology, Cellular Component
GSEA: gene set enrichment analysis
IEG: immediate-early gene
IQR: interquartile range
KS: Kolmogorov–Smirnov (statistic)
LC-MS/MS: liquid chromatography–tandem mass spectrometry
MCI: mild cognitive impairment
MS1 / MS2: full-scan (precursor) / tandem (fragment) mass spectrometry
MTG: middle temporal gyrus
NeMO: NeMO Analytics (multi-omic data exploration platform)
NIA-AA: National Institute on Aging–Alzheimer’s Association
NIA-Reagan: National Institute on Aging–Reagan Institute (criteria)
ORA: over-representation analysis
PCA: principal component analysis
PC1 / PC2: principal component 1 / principal component 2
PMI: postmortem interval
PRM-MS: parallel reaction monitoring–mass spectrometry
PV: parvalbumin
RIN: RNA integrity number
RNA-seq: RNA sequencing
SD: standard deviation
SE: standard error
TOST: two one-sided tests

## Declarations Funding

This work was supported by 5R01 AG072643 to CAB, HM, and PW. We are grateful to the Banner Sun Health Research Institute Brain and Body Donation Program of Sun City, Arizona, for the provision of human biological materials. The Brain and Body Donation Program has been supported by the National Institute of Neurological Disorders and Stroke (U24 NS072026 National Brain and Tissue Resource for Parkinson’s Disease and Related Disorders), the National Institute on Aging (P30 AG019610 and P30AG072980, Arizona Alzheimer’s Disease Center), the Arizona Department of Health Services (contract 211002, Arizona Alzheimer’s Research Center), the Arizona Biomedical Research Commission (contracts 4001, 0011, 05-901 and 1001 to the Arizona Parkinson’s Disease Consortium) and the Michael J. Fox Foundation for Parkinson’s Research. We are grateful to the SuperAging Research Initiative participants and study partners whose longitudinal participation and brain donation made this work possible.

Samples were contributed through the SuperAging Research Initiative. We are also grateful to the participants of the University of California, Irvine 90+ Study, which was supported by NIH grants R01AG021055, P30AG066519, and UF1AG057707. This project is supported by the following NIH Research Grants U19AG073153, R01AG045571 R56AG045571, 5R01AG067781, and by the McKnight Brain Research Foundation (MBRF). The content is solely the responsibility of the authors and does not necessarily represent the official views of the National Institutes of Health.

## Data and code availability

The mass spectrometry data from this study have been deposited to the ProteomeXchange Consortium (https://www.proteomexchange.org) via the PRIDE partner repository with the dataset identifier PXD078701 and project name "Molecular Coordination Around NPTX2 Distinguishes Cognitive Resilience from Normative Aging or Alzheimer’s Disease." Reviewers can access the dataset using the ID ’’ and the password ’XumOtH6M3VtB’.

The normalized, log-transformed data are available on [NeMO Analytics](https://nemoanalytics.org/landing/neurodegeneration/). Raw sequence data are published in Piras et al 2025 [68]. Sample metadata are provided in the accompanying supplementary file. Code for all analyses and visualizations is available at [GitHub](https://github.com/carlocolantuoni/LaoColantuoniAD).

## Author contributions

YL led the project, including study design, data analysis, interpretation, and drafting of the manuscript. MFX and SJ contributed to data analysis and interpretation. ISP, AB, SS, AA, JSl, AT, and MJH generated and processed the bulk RNA-sequencing data. KK and CHN performed the targeted PRM-MS proteomics. CG, EJR, CHK, MMC, GES, TGB, and JCT provided postmortem brain tissue and associated clinical and neuropathological data through their respective cohorts (the SuperAging Research Initiative, the UCI 90+ Study, the Banner Sun Health Brain and Body Donation Program, and the Johns Hopkins brain collection). CAB contributed to study oversight and interpretation. PFW and CC conceived and supervised the study, secured funding, and contributed to data interpretation and manuscript writing. All authors read and approved of the final manuscript.

## Competing interests

The authors declare that they have no competing interests.

---

**Supplementary Figure 1.**
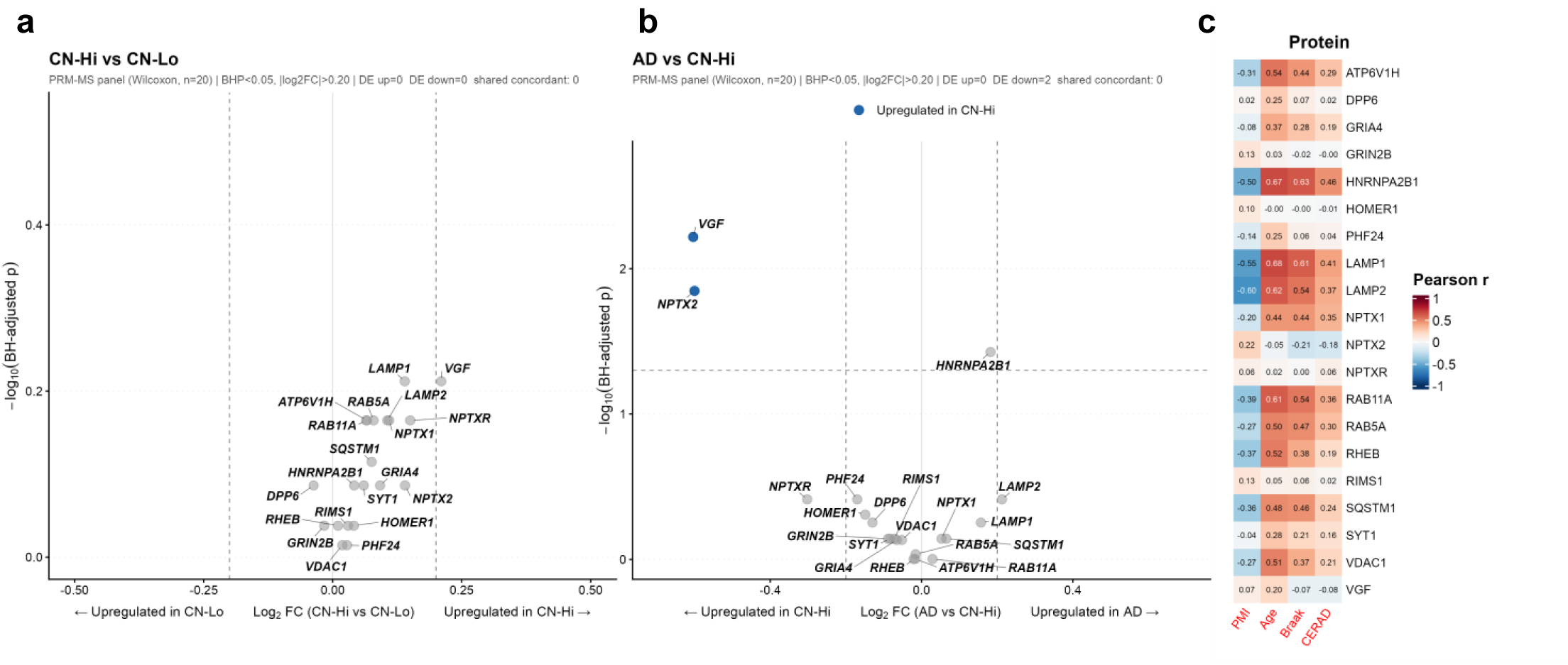
Differential expression of the 20 PRM-MS proteins. **a**, Volcano plot of differential protein abundance between CN-Hi and CN-Lo across the 20-protein PRM-MS panel (CN-Lo, n = 36; CN-Hi, n = 21 samples); positive log₂ fold change indicates higher abundance in CN-Hi. Significance was assessed by two-sided Wilcoxon rank-sum test per protein with Benjamini–Hochberg (BH) correction across the 20 proteins; the y axis shows −log₁₀(BH-adjusted p). Proteins meeting BH-adjusted p < 0.05 and |log₂ FC| > 0.20 (dashed guides) are colored by direction and labelled. **b**, As in **a**, for AD versus CN-Hi (CN-Hi, n = 21; AD, n = 38 samples); positive log₂ fold change indicates higher abundance in AD. **c**, Pearson correlation between each of the 20 PRM-MS proteins and clinical covariates (age, post-mortem interval, CERAD score, Braak stage), computed in the 135-sample PRM-MS subset and displayed as a heatmap of the correlation coefficient r (two-sided Pearson correlation; n = 135 samples).

**Supplementary Figure 2.**
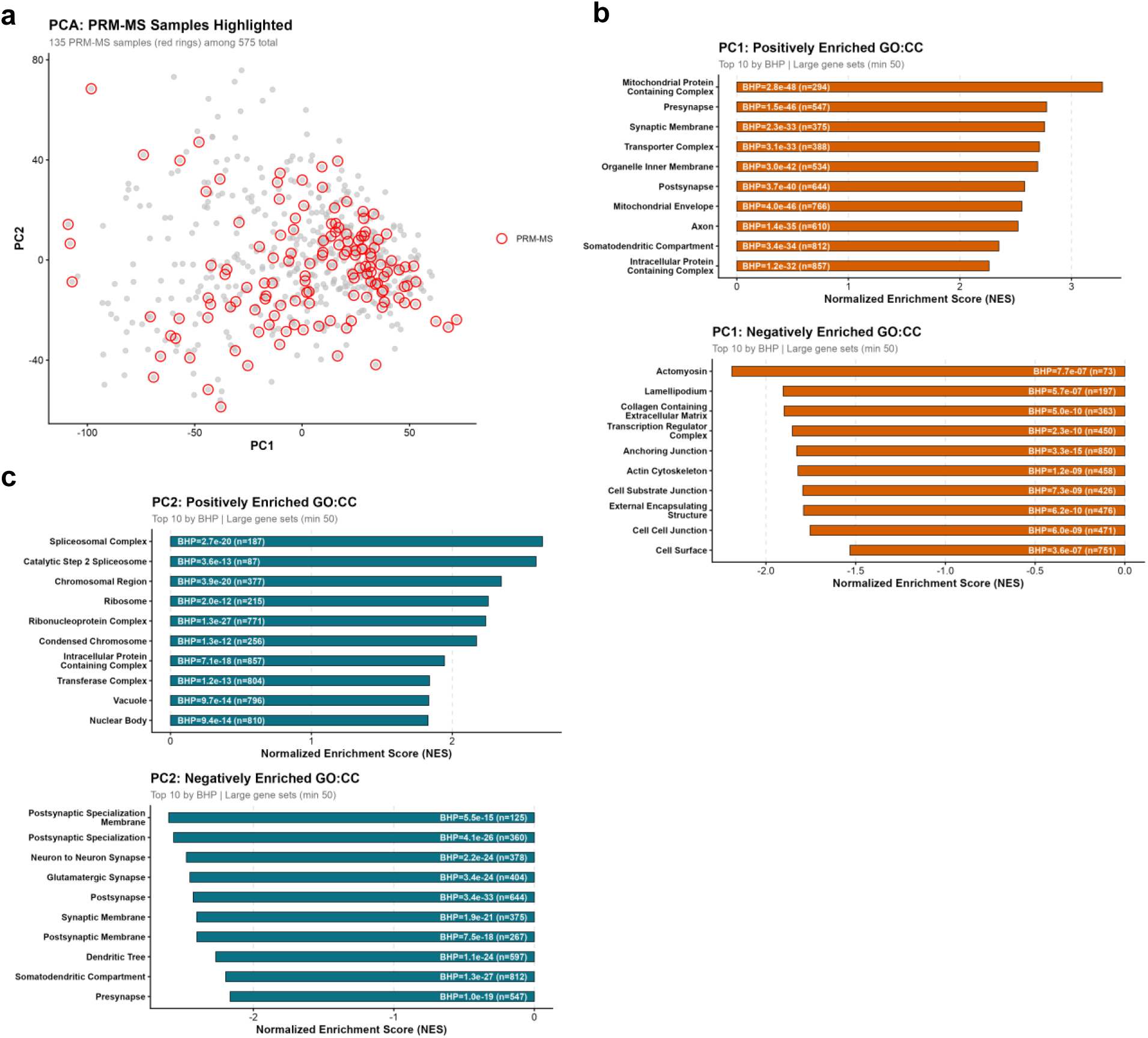
Transcriptomic PCA. **a**, PRM-MS subset membership in the principal-component space of the full cohort. All 575 samples are plotted by their precomputed PC1 and PC2 scores; the 135 samples in the PRM-MS subset are marked with hollow red rings. No statistical analysis. **b**, Gene-set enrichment of PC1 gene loadings against Gene Ontology Cellular Component (GO:CC) gene sets. Per-gene loadings were defined as the Pearson correlation between each gene and the PC1 score across all 575 samples, and tested by fast gene-set enrichment analysis (fgsea; gene sets restricted to 50–1,000 genes). Compact bar plots show the top 10 positively and top 10 negatively enriched terms ranked by BH-adjusted p-value; bar length is the normalized enrichment score (NES), and each bar is labelled in-figure with its BH-adjusted p-value (BHP) and gene-set size (n). **c**, As in **b**, for PC2 gene loadings. NES, normalized enrichment score; GO:CC, Gene Ontology Cellular Component; BHP, Benjamini–Hochberg-adjusted p-value.

**Supplementary Figure 3.**
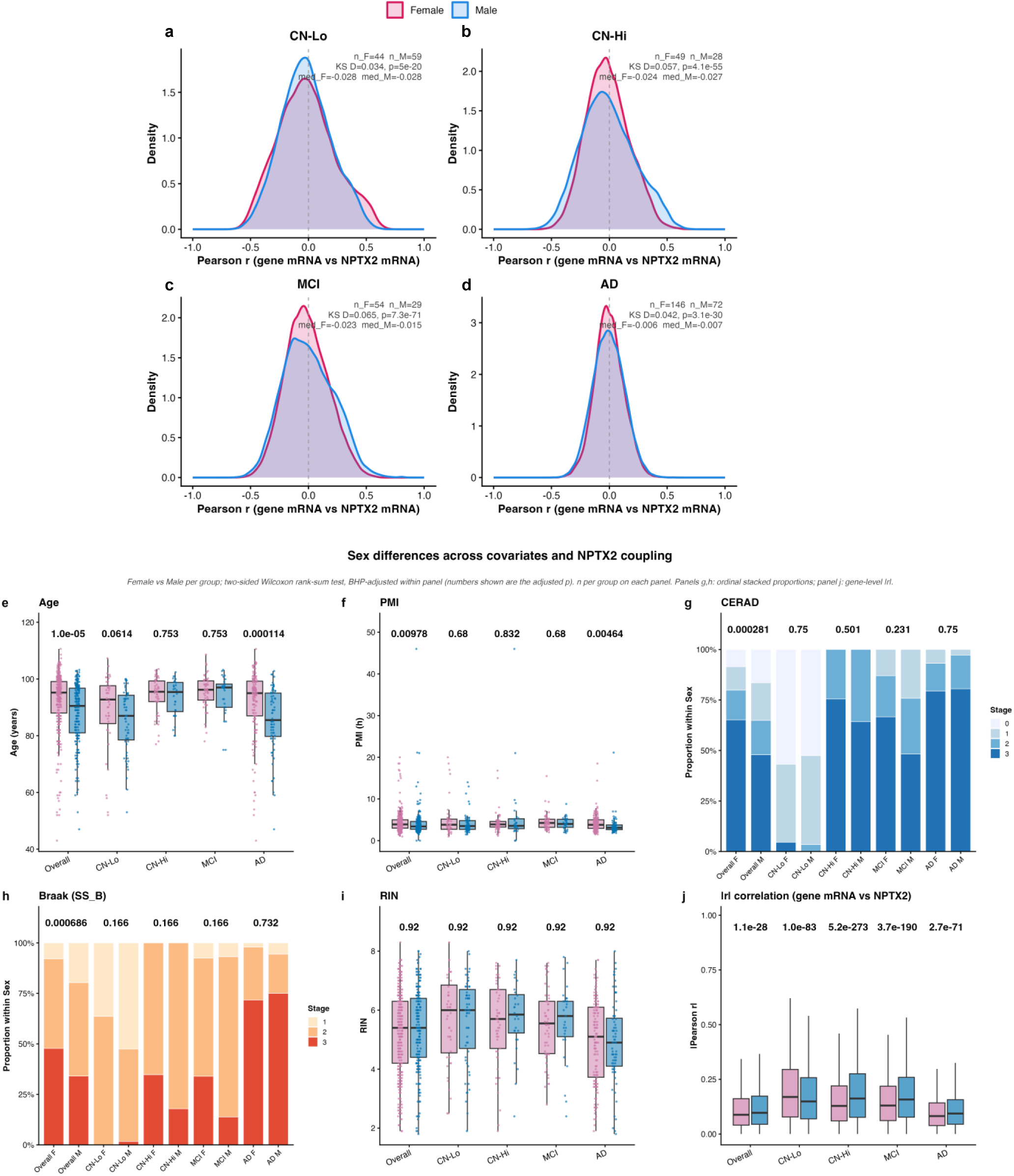
Sex differences in NPTX2 coupling and clinical covariates. **a**–**d**, Distributions of per-gene Pearson correlation with NPTX2 mRNA, compared between female and male samples within each group and shown as kernel density curves: **a**, CN-Lo; **b**, CN-Hi; **c**, MCI; **d**, AD. Each gene contributes one coefficient per sex. Female versus male distributions were compared by two-sided two-sample Kolmogorov–Smirnov (KS) test; the KS D statistic, its p-value, the per-sex sample counts (n_F, n_M) and the per-sex median r are annotated in each panel. Sample composition: CN-Lo (44 F, 59 M), CN-Hi (49 F, 28 M), MCI (54 F, 29 M), AD (146 F, 72 M). **e**–**j**, Female versus male comparison of clinical covariates and NPTX2 coupling, shown per group (Overall and the four study groups): **e**, age; **f**, post-mortem interval (PMI); **g**, CERAD score; **h**, Braak stage; **i**, RIN; **j**, gene-level |Pearson r| (gene mRNA versus NPTX2 mRNA). Panels **e**, **f**, **i** and **j** are box plots (centre line, median; box, interquartile range; whiskers, 1.5× IQR) with individual samples overlaid; panels **g** and **h** show the proportion of each ordinal stage within sex as stacked bars. All comparisons used a two-sided Wilcoxon rank-sum test, Benjamini–Hochberg-adjusted within each panel; the adjusted p-value is shown above each group. Sample composition is as in **a**–**d** (Overall, 293 F, 188 M). KS, Kolmogorov–Smirnov; PMI, post-mortem interval.

**Supplementary Figure 4.**
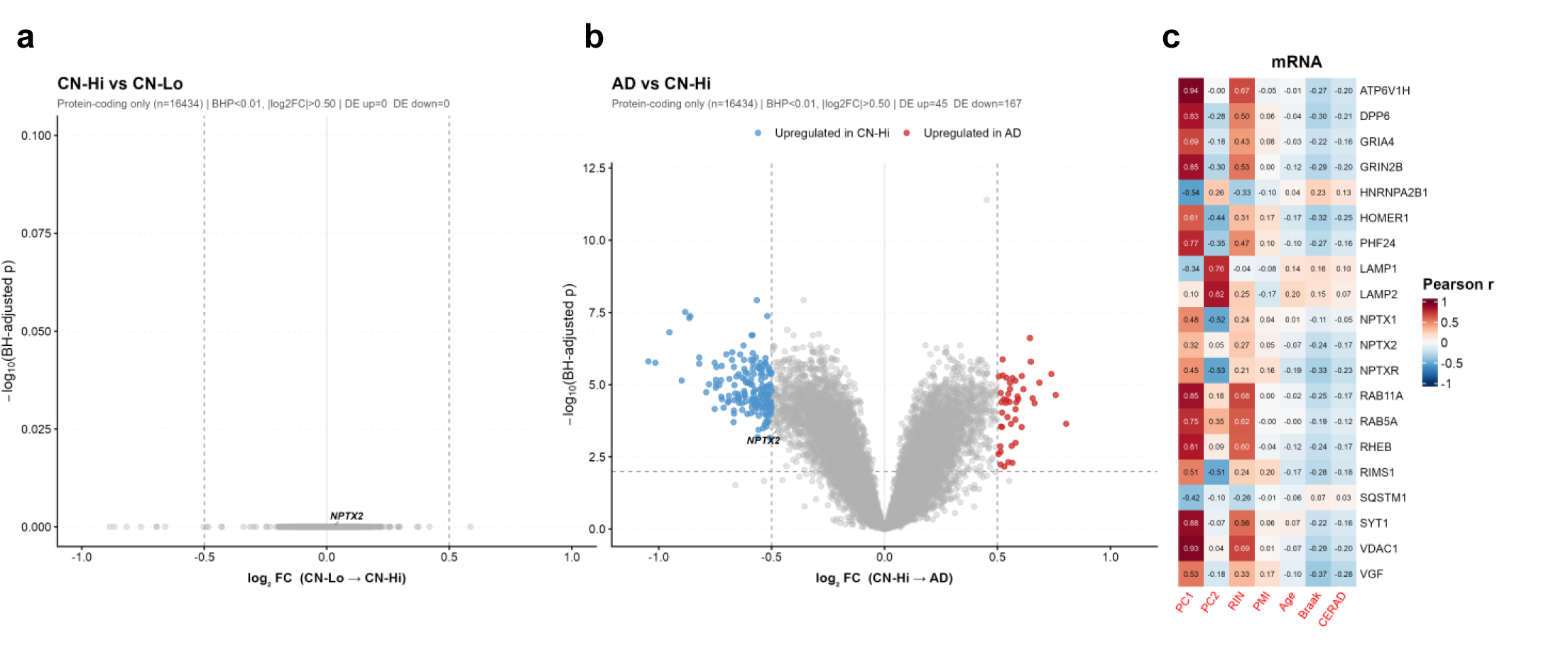
Differential expression of the Transcriptome. **a**, Volcano plot of differential expression between CN-Hi and CN-Lo across protein-coding transcripts (CN-Lo, n = 103; CN-Hi, n = 77 samples); positive log₂ fold change indicates higher expression in CN-Hi. Significance was assessed with limma moderated t-statistics (empirical Bayes, trend = FALSE, robust = TRUE) and Benjamini–Hochberg (BH) correction; the y axis shows −log₁₀(BH-adjusted p). Transcripts meeting BH-adjusted p < 0.01 and |log₂ FC| > 0.50 (dashed guides) are coloured by direction; NPTX2 is labelled. **b**, As in **a**, for AD versus CN-Hi (CN-Hi, n = 77; AD, n = 218 samples); positive log₂ fold change indicates higher expression in AD. **c**, Pearson correlation between each of the 20 targeted mRNAs and sample covariates (PC1, PC2, RIN, PMI, age, Braak stage, CERAD score), computed across all 575 samples and displayed as a heatmap of the correlation coefficient r (two-sided Pearson correlation; n = 575 samples).

**Supplementary Figure 5.**
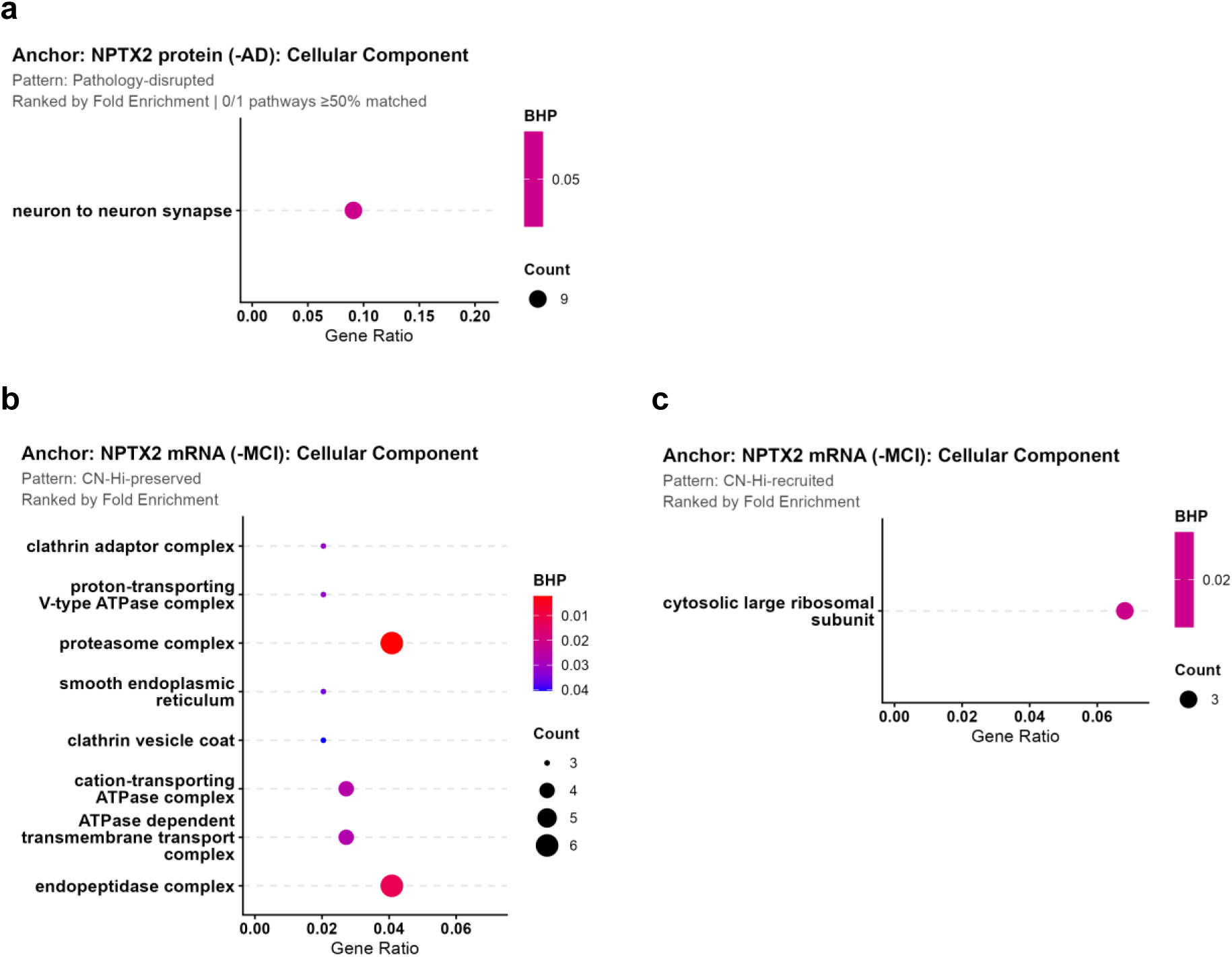
Over-representation analyses relocated from Figs 2 and 3. **a**, Over-representation analysis (Gene Ontology Cellular Component, GO:CC) of the Pathology-disrupted class from the NPTX2-protein-anchored classification. Enrichment was tested by one-sided hypergeometric test (clusterProfiler enrichGO) with Benjamini–Hochberg (BH) correction. Dot position shows the gene ratio, dot size the number of hit genes, and dot colour the BH-adjusted p-value (BHP); pathway labels in red have ≥50% of their hit genes in the pattern-matched set. **b**, As in **a**, for the CN-Hi-preserved class from the NPTX2-mRNA-anchored classification. **c**, As in **a**, for the CN-Hi-recruited class from the NPTX2-mRNA-anchored classification. GO:CC, Gene Ontology Cellular Component; BHP, Benjamini–Hochberg-adjusted p-value.

**Supplementary Table 1.**
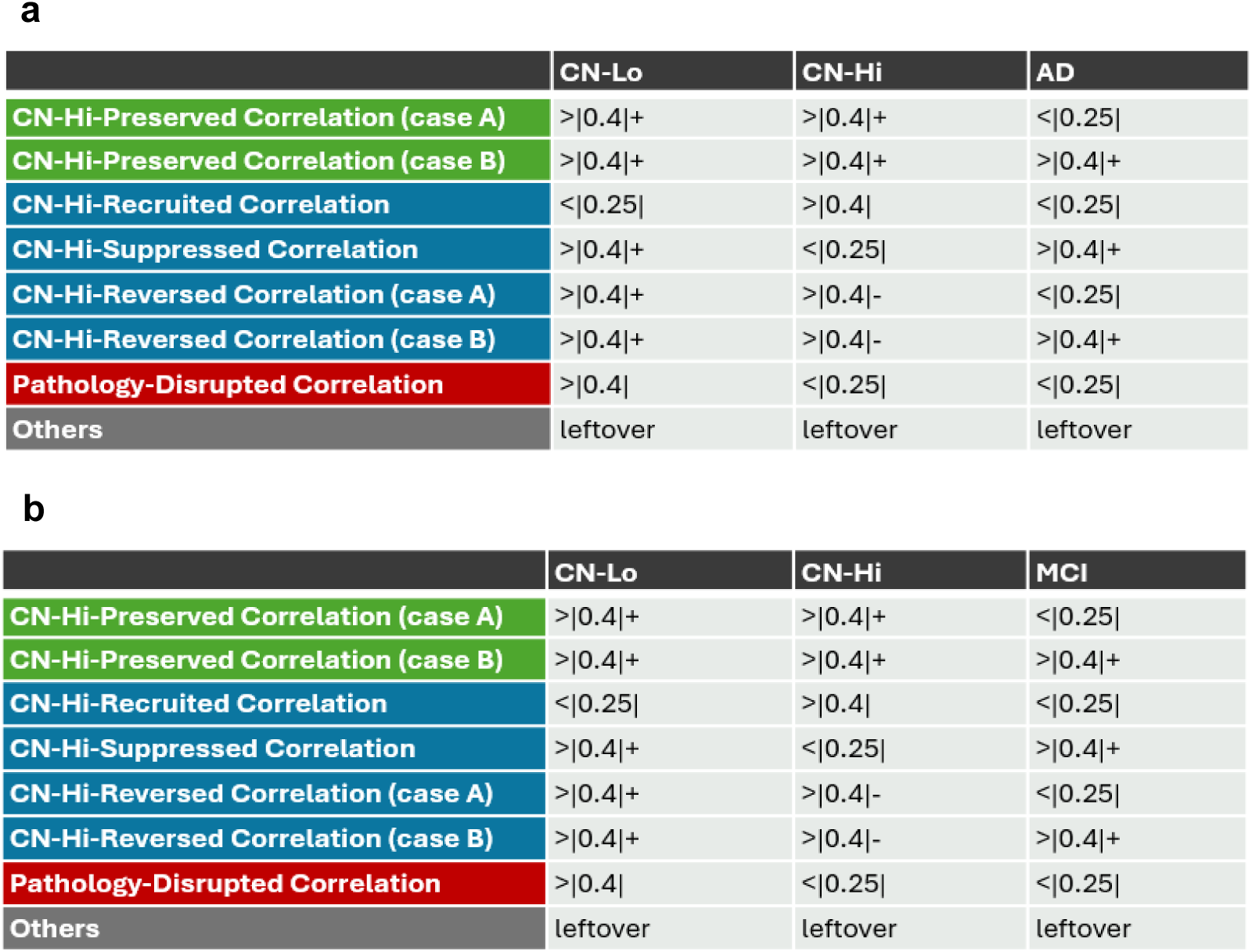
Classification rules for NPTX2 correlation trajectory patterns. **a,** AD-based classification. Each row defines one of five mutually exclusive patterns based on the direction and magnitude of Pearson correlation between a candidate gene and NPTX2 within three diagnostic groups (CN-Lo, CN-Hi, AD). Thresholds: strong, |r| > 0.40; weak, |r| < 0.25. The "+" and "−" signs denote concordant or discordant direction. Genes not meeting any pattern are classified as "Others". **b,** MCI-based classification: identical rules with MCI replacing AD. Genes assigned the same pattern and sign under both rules are flagged as trajectory-matched.

|  | CN-Lo | CN-Hi | AD |
| --- | --- | --- | --- |
| CN-Hi-Preserved Correlation (case A) | >\|0.4\|+ | >\|0.4\|+ | <\|0.25\| |
| CN-Hi-Preserved Correlation (case B) | >\|0.4\|+ | >\|0.4\|+ | >\|0.4\|+ |
| CN-Hi-Recruited Correlation | <\|0.25\| | >\|0.4\| | <\|0.25\| |
| CN-Hi-Suppressed Correlation | >\|0.4\|+ | <\|0.25\| | >\|0.4\|+ |
| CN-Hi-Reversed Correlation (case A) | >\|0.4\|+ | >\|0.4\|- | <\|0.25\| |
| CN-Hi-Reversed Correlation (case B) | >\|0.4\|+ | >\|0.4\|- | >\|0.4\|+ |
| Pathology-Disrupted Correlation | >\|0.4\| | <\|0.25\| | <\|0.25\| |
| Others | leftover | leftover | leftover |

**Supplementary Table 1. Classification rules for NPTX2 correlation trajectory patterns.**
|  | CN-Lo | CN-Hi | MCI |
| --- | --- | --- | --- |
| CN-Hi-Preserved Correlation (case A) | >\|0.4\|+ | >\|0.4\|+ | <\|0.25\| |
| CN-Hi-Preserved Correlation (case B) | >\|0.4\|+ | >\|0.4\|+ | >\|0.4\|+ |
| CN-Hi-Recruited Correlation | <\|0.25\| | >\|0.4\| | <\|0.25\| |
| CN-Hi-Suppressed Correlation | >\|0.4\|+ | <\|0.25\| | >\|0.4\|+ |
| CN-Hi-Reversed Correlation (case A) | >\|0.4\|+ | >\|0.4\|- | <\|0.25\| |
| CN-Hi-Reversed Correlation (case B) | >\|0.4\|+ | >\|0.4\|- | >\|0.4\|+ |
| Pathology-Disrupted Correlation | >\|0.4\| | <\|0.25\| | <\|0.25\| |
| Others | leftover | leftover | leftover |

## Notes

### Competing Interest Statement

The authors have declared no competing interest.

### Summary of Updates

In this revision, we split controls into CN-Lo (low pathology controls) and CN-Hi (high pathology controls), based on CERAD and Braak stages.

